# Microbiome time series data reveal predictable patterns of change

**DOI:** 10.1101/2023.06.08.544023

**Authors:** Zuzanna Karwowska, Paweł Szczerbiak, Tomasz Kosciolek

## Abstract

**Background:** The gut microbiome is crucial for human health and disease. Longitudinal studies are gaining importance in understanding its dynamics over time, compared to cross-sectional approaches. Investigating the temporal dynamics of the microbiome, including individual bacterial species and clusters, is essential for comprehending its functionality and impact on health. This knowledge has implications for targeted therapeutic strategies, such as personalized diets and probiotic therapy.

**Results:** Here, by adopting a rigorous statistical approach, we aim to shed light on the temporal changes in the gut microbiome and unravel its intricate behavior over time. We leveraged four long and dense time series of the gut microbiome in generally healthy individuals examining how its composition evolves as a community and how individual bacterial species behave over time. We also explore whether specific clusters of bacteria exhibit similar fluctuations, which could provide insights into potential functional relationships and interactions within the microbiome Our study reveals that despite its high volatility, the human gut microbiome is stable in time and can be predicted based solely on its previous states. We characterize the unique temporal behavior of individual bacterial species and identify distinct longitudinal regimes in which bacteria exhibit specific patterns of behavior. Finally, through cluster analysis, we identify groups of bacteria that exhibit coordinated fluctuations over time.

**Conclusions:** Our findings contribute to our understanding of the dynamic nature of the gut microbiome and its potential implications for human health. The provided guidelines support scientists studying gut microbiome complex dynamics, promoting further research and advancements in microbiome analysis.

## Introduction

The gut microbiome plays a crucial role in human health and disease. The majority of microbiome studies have relied on cross-sectional data, providing only snapshots of its composition at a specific time point^1–3^. In recent years, there has been a growing recognition of the importance of longitudinal studies, which enable us to explore the dynamics of the gut microbiome over time^4–10^.

Longitudinal studies, especially those involving dietary or antibiotic interventions, have revealed that while the gut microbiome remains relatively stable over time, it also exhibits considerable variability in response to perturbations^11–14^. Therefore, investigating the temporal dynamics of the gut microbiome, including the behavior of different bacterial species over time and the identification of bacterial clusters exhibiting similar fluctuations, is critical for comprehending its functionality and impact on human health. Understanding the dynamics of the gut microbiome, including its temporal changes and bacterial relationships, enhances our comprehension of its role in health and disease. This knowledge has significant implications for the development of targeted therapeutic strategies, such as personalized diet, probiotic therapy, and fecal microbiota transplantation (FMT), enabling personalized interventions^15,16^. By examining the temporal changes in the gut microbiome, we can identify disease-associated patterns and trends, facilitating the development of interventions to restore or modulate the microbiome for improved health outcomes. Furthermore, studying the behavior of individual bacterial species over time allows us to pinpoint key contributors to dysbiosis and target them with interventions like probiotics to restore microbial balance. By identifying the beneficial bacteria crucial for human health, we can establish a comprehensive health index that considers diverse physiological and microbial parameters. This approach allows us to proactively monitor the microbiome’s composition and detect subtle changes in the overall health status. Through this proactive monitoring, we can intervene in a timely manner, leveraging targeted probiotics and other interventions to restore microbial balance and prevent the onset or progression of diseases^17^. In summary, comprehending the dynamics of the gut microbiome enables the development of more precise and effective therapeutic strategies, ultimately leading to improved human health and the prevention or treatment of microbiome-associated disorders.

Here, by adopting a longitudinal approach, we aim to shed light on the temporal changes in the gut microbiome and unravel its intricate behavior over time. In this study, we investigate the temporal dynamics of the gut microbiome, examining how its composition evolves as a community and how individual bacterial species behave over time. We also explore whether specific clusters of bacteria exhibit similar fluctuations, which could provide insights into potential functional relationships and interactions within the microbiome.

In contrast to the prevailing use of observational approaches in many previous studies^7,18^, our research distinguishes itself by employing statistical methods to analyze human gut microbiome time series data. This distinction is significant as statistical analysis allows for a more rigorous examination of microbial dynamics, enabling the identification of patterns, trends, and associations that may have been overlooked. By applying statistical tests, we not only confirm the consistency of our results with prior findings but also provide a systematic and reproducible framework that quantifies the behaviors of individual bacterial species. This quantitative approach adds depth and reliability to our understanding of the gut microbiome and opens up new avenues for personalized medicine and targeted interventions.

## Results

In the first part we describe the behavior of the microbiome over time as a whole. We examine whether it exhibits white noise behavior, is stationary, and seasonal, and whether we can predict its change over time. We also demonstrate how the taxonomy changes over time. The second part focuses on the analysis of individual amplicon sequence variants (ASV) that constitute the microbiome and their behavior over time. We present a methodology through which we describe each ASV using a longitudinal feature vector. Furthermore, we demonstrate the existence of groups of ASVs that exhibit similar behavior over time. Finally, in the third part, we present the results of graph analysis, which show groups of bacteria that fluctuate together over time.

### Whole community analysis

For all analyses in this study, we used two publicly available 16S rRNA marker gene sequencing datasets containing data on human gut microbiome from four, adult individuals with no reported diseases (Table 1). We maintain consistent nomenclature by referring to each individual using the names originally assigned to them in the original studies, that is male and female subjects for the first dataset and donorA and donorB for the second dataset.

**Table 1.** Dataset summary.

| Study | Subject ID | Time points | ASV | Additional information |
| --- | --- | --- | --- | --- |
| Moving pictures of the human microbiome<sup>7</sup> | male | 443 | 1 253 | subjects were undergoing antibiotic treatment prior to sampling |
|  | female | 185 | 551 |  |
| Host lifestyle affects human microbiota on daily timescales<sup>8</sup> | donorA | 365 | 1 524 | subject was traveling in days 70 to 122, which caused a change in gut microbiome composition |
|  | donorB | 252 | 1 569 | subject suffered from food poisoning between day 150 to 159 |

#### The human gut microbiome is individual but stable over time

Initially, we assessed the temporal dynamics of the human gut microbiome. PCoA analysis revealed distinct clusters for each subject’s microbiome, indicating host-specificity and dynamic changes over time (Fig. 1A and C, Supplementary Fig. 1). To explore the collective dynamics, we computed alpha diversity indices (Shannon diversity index and Faith’s Phylogenetic Diversity index) and observed fluctuations but consistent mean values (Fig. 1B Supplementary Fig. 2). For donorA, linear regression analysis demonstrated a trend towards baseline following perturbations (yellow box in the right panel of Fig. 1B) despite day-to-day variations.

**Figure 1.**
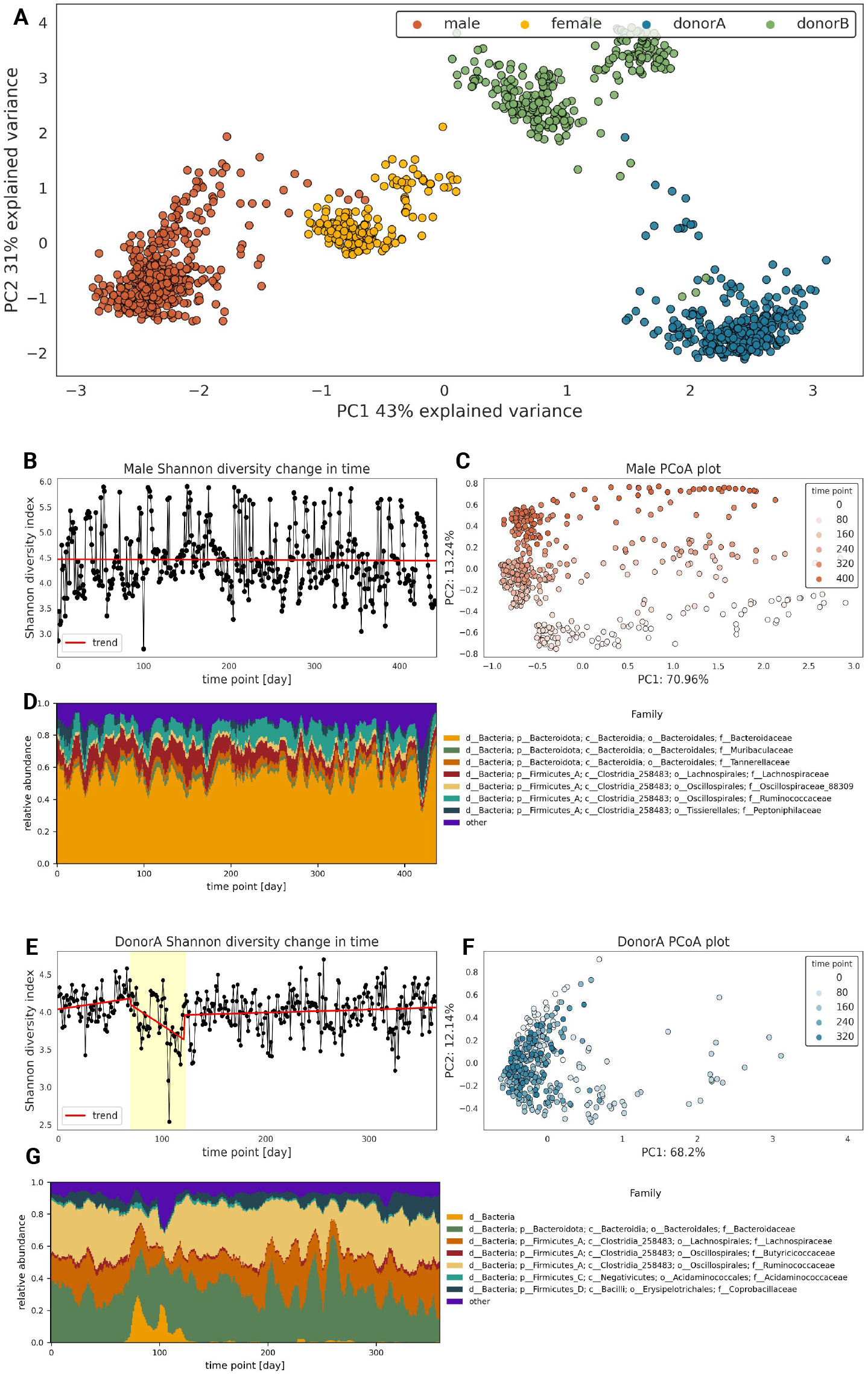
Analysis of general behavior of gut microbiome in time. **A**. The scatterplot of the two first coordinates of Principal Coordinates Analysis (PCoA) on Aitchinson distance. **B**. Lineplot of male subject Shannon diversity index over time. C. Scatterplot of the two first coordinates of male subject PCoA on Aitchinson distance matrix. **D**. Rolling mean of male subject taxonomy composition in time (window=14days). **E**. Lineplot of donorA Shannon diversity index over time. The yellow box showcases days 70 to 122 when the subject was traveling. **F**. Scatterplot of the two first coordinates of donorA PCoA on Aitchinson distance matrix. **G**. Rolling mean of DonorA subject taxonomy composition in time (window=7days).

**Figure 2.**
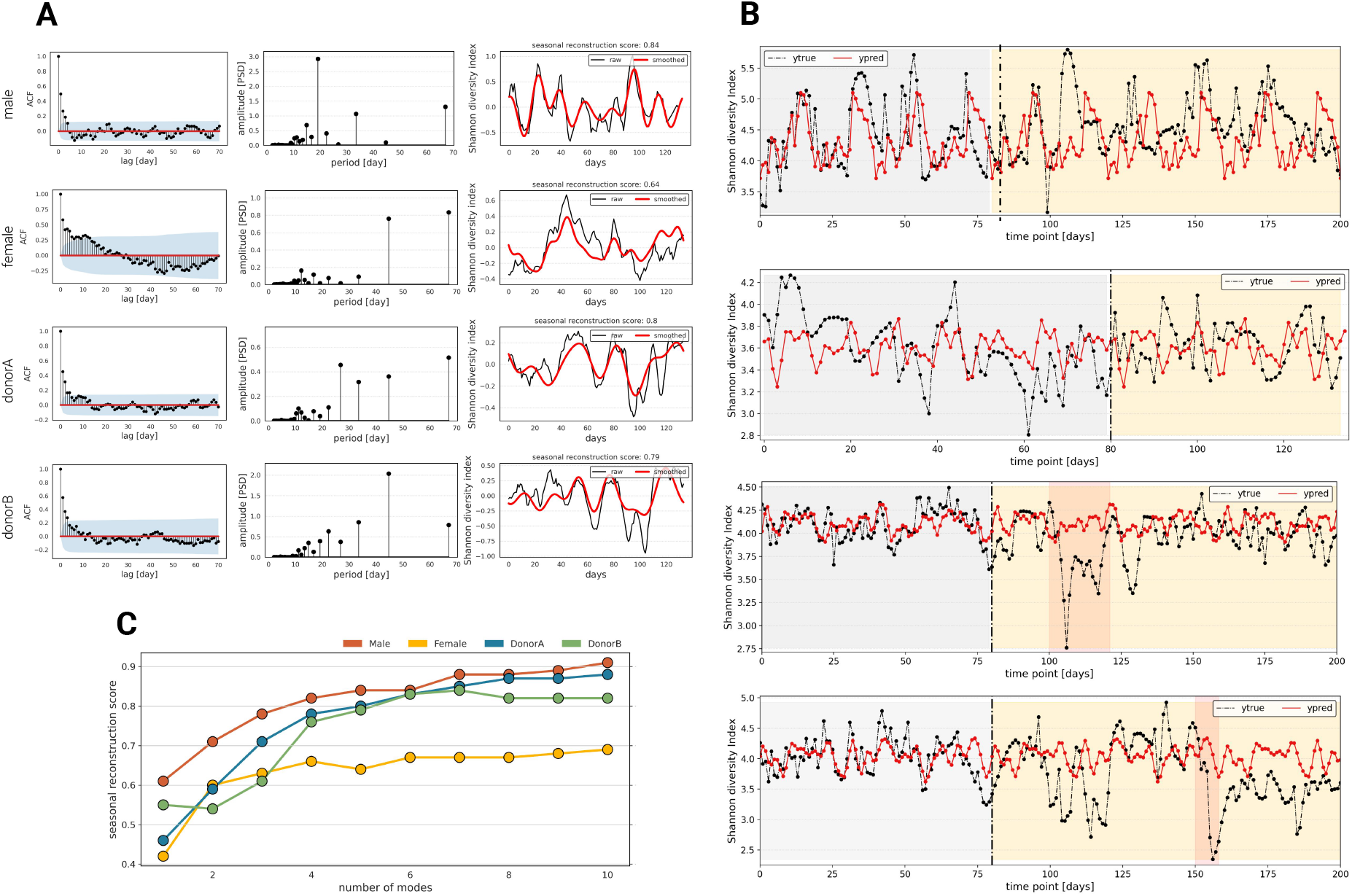
Shannon diversity index behavior in time. **A**. Left: autocorrelation coefficient plots (blue area shows the significance level); middle: spectrograms showing most dominant seasonalities of the human gut microbiome; right: reconstruction of alpha diversity using 5 dominant seasonalities plotted against raw alpha diversity change in time. **B**. Prediction of alpha diversity change in time using a dynamic ARIMAX model with FFT seasonalities. **C**. Plot showing the relationship between a number of used seasonalities to reconstruct alpha diversity and the seasonal reconstruction score.

Analyzing longitudinal human gut microbiome data from four individuals, we found that while each person has a unique microbiome composition, there are common dominant bacterial taxa (Fig. 1D, Supplementary Fig. 1). We observed a shared trend where certain bacterial taxa consistently maintained high abundance throughout the duration of the longitudinal data, while others showed lower abundance and temporal variability, appearing intermittently over time^18,19^. Specifically, *Ruminococcaceae, Lachnospiracea, Bacteroidaceae, Oscillospiraceae 88309*, and *Acidaminococcaceae* families dominate in all four subjects. These bacterial families are commonly found in the human gut microbiome and have a shared ability to ferment dietary fibers and produce short-chain fatty acids. They contribute to gut health by providing energy to gut cells and exhibiting metabolic versatility^20–23^.

#### Predictability of human gut microbiome

In targeted microbiome therapy, the goal is to anticipate the response of the gut microbiome community following the administration of therapy^24^. To this end, we aimed to investigate whether the gut microbiome exhibits properties of a predictable time series or if it behaves as a white noise process.

Given the high-dimensional nature of gut microbiome data, the first objective of this study was to investigate the behavior of the human gut microbiome as a unified entity. To achieve this, we employed alpha diversity as a means of biologically reducing the dimensionality of the data to a univariate parameter. Here, we computed two diversity indices, namely the Shannon diversity index and Faith’s phylogenetic diversity index, to quantitatively assess the diversity and evolutionary relationships among the microbial taxa present in the human gut (Supplementary Fig. 2). Finally, we tested each time series for characteristics such as the similarity to the white noise process, stationarity, and the presence of seasonality in the data (Fig. 2).

We investigated whether the human gut microbiome exhibits white noise behavior by analyzing Shannon’s diversity index and Faith’s phylogenetic diversity index of each subject’s gut microbiome time series. Autocorrelation and partial autocorrelation analysis, as well as, Ljung-Box tests on 30 lags revealed the presence of both autocorrelation and partial autocorrelation, indicating gut microbiome dependence on its previous states (Fig. 2A (left panels), Supplementary Fig. 3, Supplementary Table 1). The low flatness score suggested the predictability of the gut microbiome as a whole. Unit root tests confirmed the stationary nature of the human microbiome, indicating a relatively constant composition over time. Volatility clustering analysis identified regions of increased variance (Supplementary Fig. 4). However, the lack of metadata limited the understanding of the underlying factors driving this variability. The observed characteristics support the predictability of the gut microbiome’s future behavior in general, but local perturbations require relevant metadata for a comprehensive understanding.

**Figure 3.**
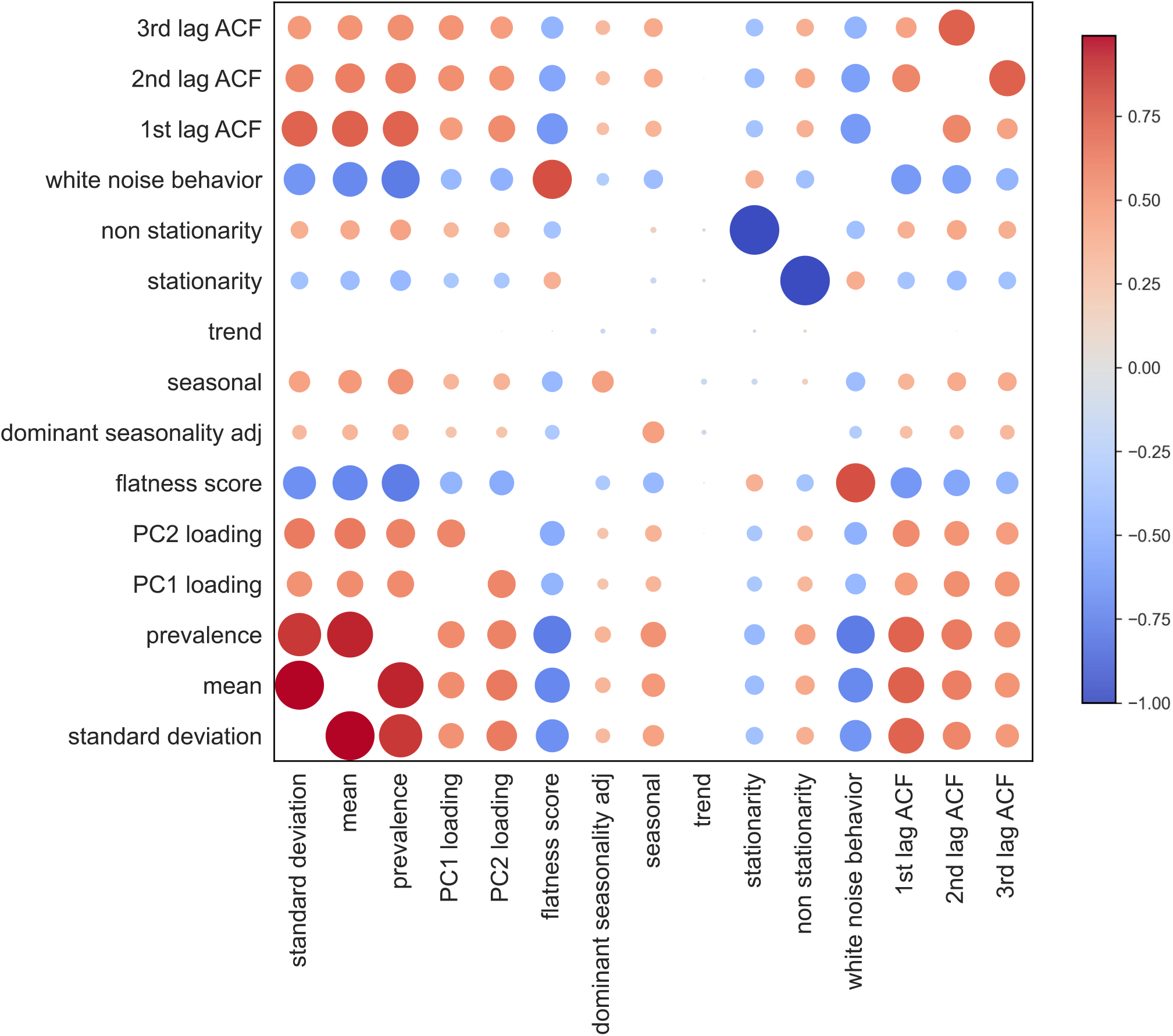
Correlation of longitudinal characteristics of human gut microbiome. Spearman correlation matrix, representing the relationships between different longitudinal behaviors of the human gut microbiome.

**Figure 4.**
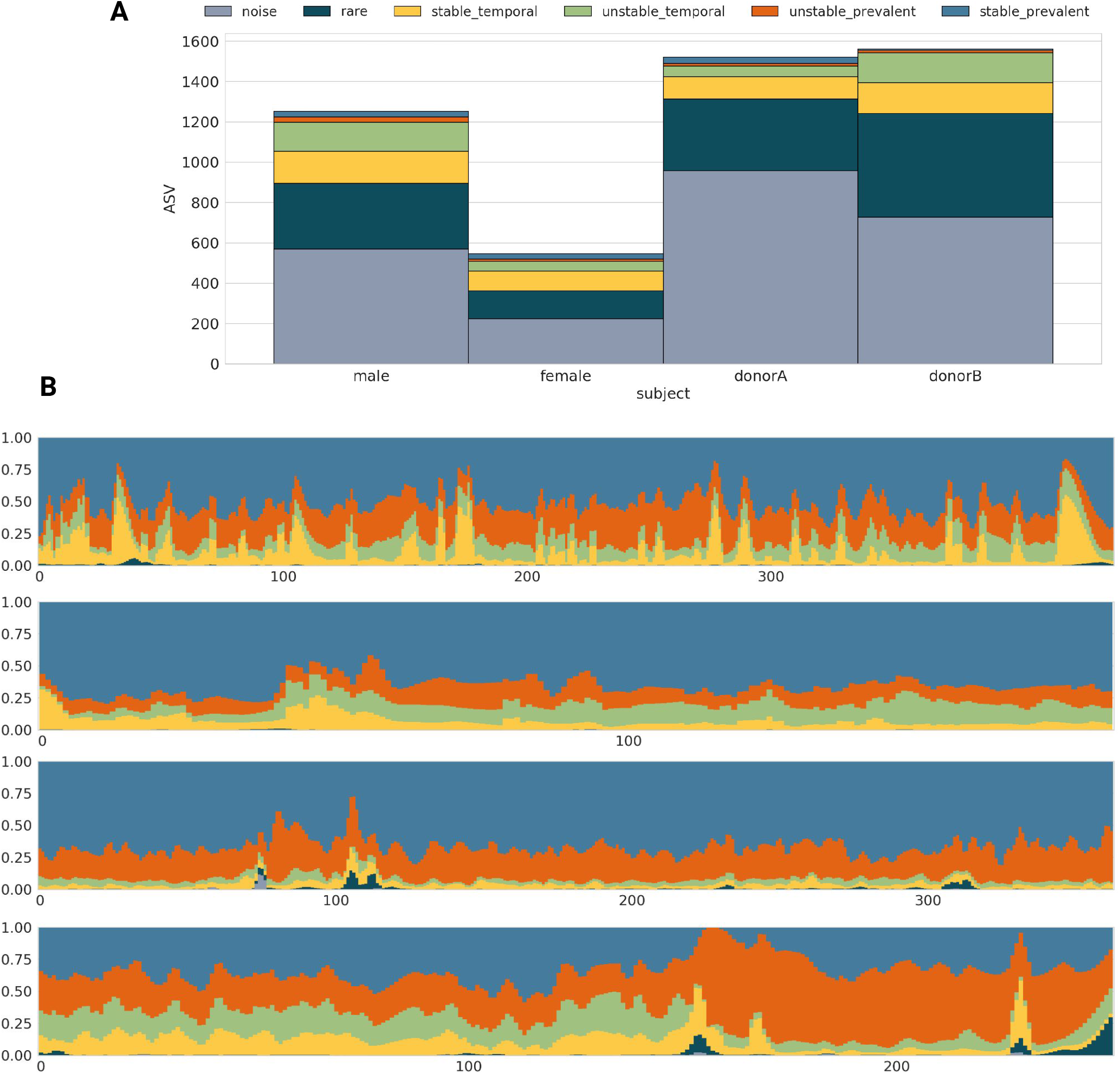
Longitudinal characteristics of human gut microbiome. **A**. Bar plots showing the number of bacteria exhibiting specific regimes in each subject. **B**. The fluctuation in counts over time for each longitudinal regime using a stacked barplot. Panels represent male, female, donor A and donorB regimes fluctuation in time. X-axis represents time points in days and y-axis represents relative abundance of particular regimes.

Using the analysis of the autocorrelation function and partial autocorrelation function analysis we perceived the presence of seasonal components in the longitudinal human microbiome data. Using Fast Fourier transform (FFT) analysis further, we detected multiple dominant seasonal patterns unique to each subject (middle panels in Fig. 2A, Supplementary Fig. 3C-D). Additionally, low power density spectra indicated the presence of noise and short artificial seasonalities. To assess the reliability of the detected seasonality, we introduced a measure called the seasonal reconstruction score. This score quantifies the Spearman correlation between the raw signal and the signal reconstructed using *N* Fourier seasonalities. To validate these patterns, we performed inverse FFT and calculated the seasonal reconstruction score, finding that at least five seasonalities were required for optimal reconstruction (Fig. 2A (right panels) and supplementary Fig. 3C-D).

Finally, we aimed to validate the predictability of the human gut microbiome using a dynamic ARIMAX model. The model incorporated dominant seasonal patterns identified through FFT analysis, as standard SARIMA models were deemed insufficient. ARIMAX models were trained on the initial 80 days of data for each subject, and the parameters were optimized through grid search cross-validation. The model provided a good fit to the training data and satisfactory performance on the test set (Fig. 2B, Supplementary Fig. 5). However, there were certain regions in the time series where the model was unable to accurately predict fluctuations, as seen in donorA between days 100 and 120, where a drop in alpha diversity occurred, and in donorB after day 150, during a period of diarrhea (Supplementary Fig. 5). Nonetheless, our primary aim was to test the self-explanatory nature of the human gut microbiome and evaluate the ability of the model to predict fluctuations in the absence of metadata.

**Figure 5.**
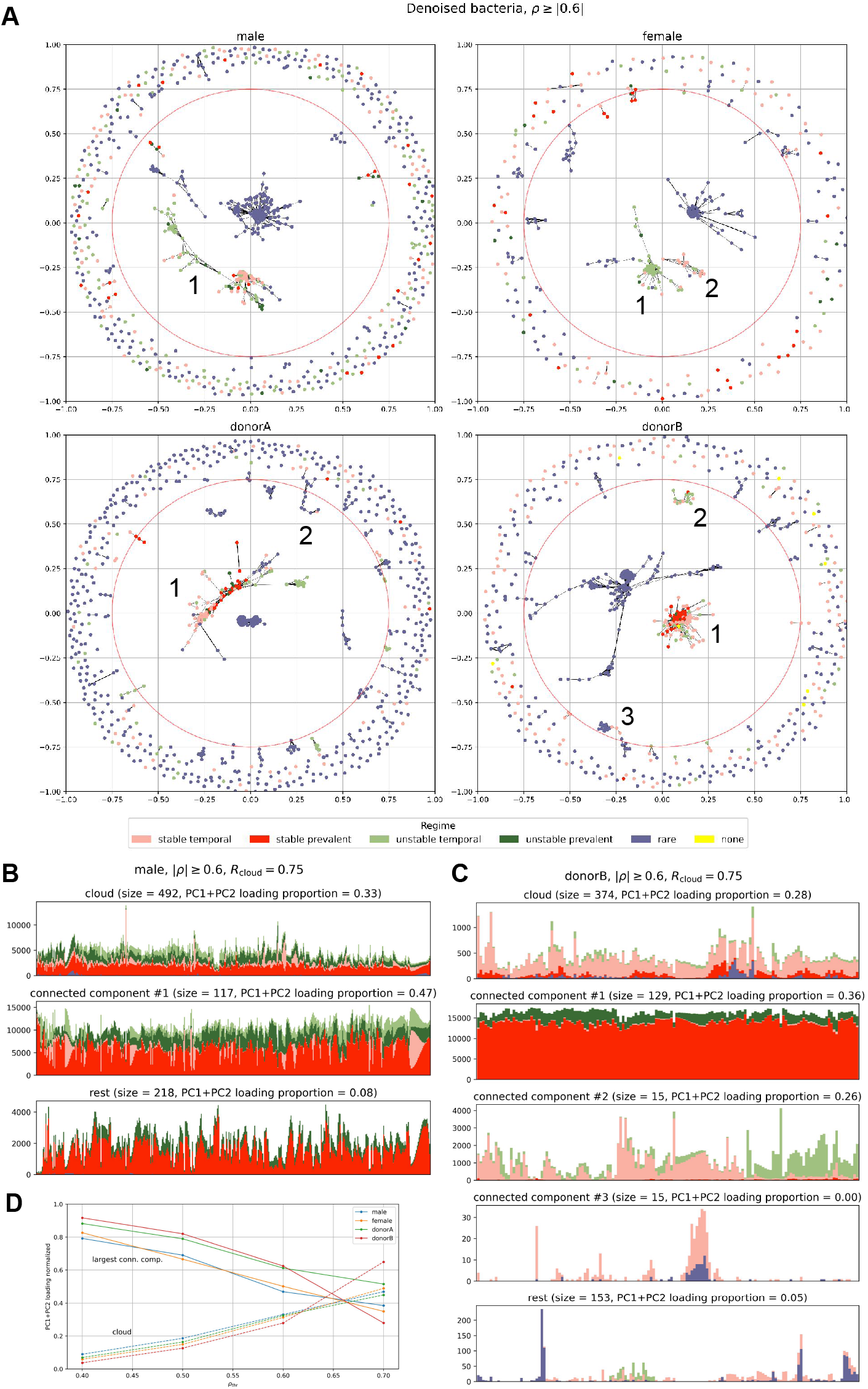
Cluster analysis performed with NetworkX. **A**. network (one panel per subject) of bacterial species (nodes) where connections (edges) represent proportionality equal or stronger than *ρ*_thr_ = 0.6 (equivalent to |*ρ*| ≥ 0.6). Colors correspond to longitudinal regimes defined in section “Individual features analysis” (“none” represents bacteria that didn’t pass rarefaction). Red circle in each panel separates the inner part from the “cloud”. **B**. change in total counts after rarefaction over time for male subjects stratified by regime (color) and group of bacteria (panels) i.e. cloud, largest connected components (numbers in the graph) and the rest. **C**. the same as in B but for donorB. *R*_cloud_ represents a diameter that separates the inner part from the cloud. **D**. PC1+PC2 loading against *ρ*_thr_ for bacteria located in the cloud and largest connected components.

### Individual features analysis

To gain a comprehensive understanding of the behavior of individual bacteria within the human gut microbiome, we generated longitudinal feature vectors for each taxon (represented by ASVs) that captured their characteristics over time. Each feature vector was of length 12 (Table 2). The vectors included general time series characteristics that is: white noise behavior, stationarity, presence of a seasonal component, and impact on the variability of the overall time series. We simplified the quantification of bacterial behavior by defining two artificial characteristics: noise and seasonal behavior, based on fixed thresholds. Bacteria exhibiting random behavior with no autocorrelation and a flatness score above 0.4 were classified as “noise” (Supplementary Fig. 6). We used Fast Fourier Transform analysis to identify dominant seasonal patterns for each taxon. Bacteria were classified as seasonal if their seasonal reconstruction score for 6 Fourier modes was at least 0.4 (Table 2, Supplementary Fig. 10, Methods section).

**Table 2.**
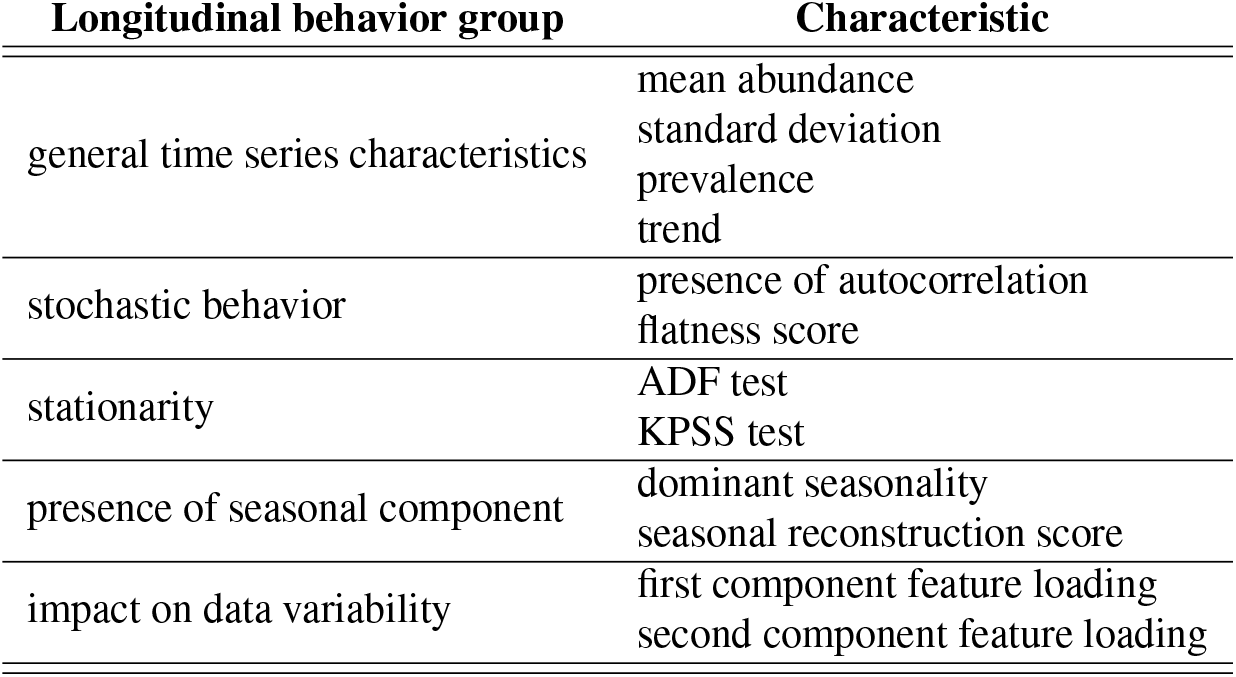
Characteristics of gut microbiome time series used to construct a feature vector.

First, we sought to identify unique longitudinal signatures in each subject’s gut microbiome by individually analyzing selected longitudinal characterictics (Table 2). The analysis of bacterial abundance in all four subjects revealed distinct patterns. A significant proportion of bacteria were classified as rare or predominantly absent, while a smaller fraction was consistently present in the gut microbiome. Another subset appeared intermittently (Supplementary Fig. 8). Surprisingly, almost half of each individual’s microbiome was classified as noise, likely resulting from technical factors and metabolic conditions^25^ (Supplementary Fig. 7). A considerable portion of ASVs in each subject displayed stationary behavior (Supplementary Fig. 9). The analysis of seasonality in the human gut microbiome revealed that only a small fraction of bacteria exhibited seasonal behavior, indicating a high variability in their patterns (Supplementary Fig. 12). Interestingly, shorter seasonalities were predominantly characterized as noise, while longer seasonalities showed higher seasonal reconstruction scores (Supplementary Fig. 11). Additionally, the unique features associated with seasonality were found to be consistent with the seasonalities derived from alpha diversity analysis, reinforcing the presence of meaningful seasonal characteristics in the gut microbiome (Supplementary Fig. 12).

#### Longitudinal regimes of the human gut microbiome

After creating a longitudinal characteristics vector for each ASV, we aimed to identify groups of taxa exhibiting similar patterns of behavior in the human gut microbiome. For this purpose, we generated a Spearman correlation matrix from feature vector variables to determine correlated characteristics (Fig. 3).

After analyzing the correlation matrix of longitudinal characteristics, we categorized bacteria into six groups based on their specific longitudinal behavior: 1) prevalent and stable, 2) prevalent but unstable, 3) temporal and stable, 4) temporal but unstable, 5) rare, and 6) noise.

The prevalent and stable group consists of bacteria that are consistently present in over 90% of the time series and exhibit stationarity. These bacteria show high abundance and are not classified as noise, suggesting their stable presence in the human gut microbiome despite environmental changes. Bacteria classified as prevalent but unstable are also present in over 90% of the time series but do not demonstrate stationarity. Due to their high abundance and non-white noise behavior, these bacteria have a significant impact on time series variance (Supplementary Fig. 14). The temporal and stable group comprises bacteria that are present in more than 20% but less than 90% of the time series, exhibit stationarity and are not classified as noise. Bacteria in the temporal but unstable group share the same prevalence patterns as the temporal and stable group but lack stationarity, indicating fluctuating behavior over time. All groups, except for the prevalent and stable group, are classified as a part of the volatile gut microbiome. We hypothesize that these bacteria have more complex metabolic requirements and are more responsive to environmental changes such as diet and medications^19,26,27^. Additionally, we identified two distinct groups: rare bacteria and noisy bacteria. Rare bacteria are present in less than 20% of the time series but do not exhibit white noise behavior. On the other hand, noisy bacteria are characterized by low abundance and occurrence, stationarity, and minimal impact on time series variance (low PCoA loading).

Next, we performed an analysis to determine the prevalence of each longitudinal group in every subject. We observed that in all four subjects, bacteria categorized as rare or noisy accounted for over 50% of all rarefied ASVs (Fig. 4A) or even 70-90% when raw counts are considered (Supplementary Table 2). The next, most abundant group were the temporal bacteria (both stable and unstable). Moreover, every subject contained a small portion of prevalent and stable bacteria (Fig. 4A). Finally, for all subjects, we analyzed the temporal fluctuations of each longitudinal group over time. Interestingly, although the stable bacteria group representatives were less numerous, they constituted the majority of the time series in terms of abundance. This was followed by a smaller fraction of temporal bacteria exhibiting higher volatility. Additionally, rare bacteria appeared for short periods of time, and bacteria defined as noise, despite being the most numerous group, were nearly undetectable when taking into account their abundance (Fig. 4B).

### Analysis of bacterial clusters

In order to shed more light on bacterial redundancy (i.e. how many bacteria behave similarly) and their relationships we performed cluster analysis. First, for each subject, we computed proportionality (which is a recommended method for correlation analysis of compositional data^28^. The proportionality matrix *ρ* (of shape *N × N*, where *N* is the number of all species in the dataset) has been transformed to pseudo-distance matrix as *D* = 1− | *ρ* | (meaning that any two species that correlate or anti-correlate have distance 0 and distance 1 if they don’t). Next, we generated NetworkX graphs using spring layout (see details in Methods section) - results are presented in Fig. 5.

We identified three distinct regions in the network: a large cluster in the center of the graph that consists of stochastic and very rare taxa that didn’t pass the rarefaction step (visible only when both noisy and signal species are considered - see Supplementary Fig. 15A), medium-size connected components and a distant cloud of bacteria comprising mostly singletons i.e. species that do not (anti)co-occur with the others. Fig. 5A shows denoised bacteria (i.e. species after rarefaction that don’t behave as white noise) colored by the longitudinal regime for *ρ*_thr_ = 0.6 i.e. | *ρ* | ≥ 0.6 (see Supplementary Fig. 15L–M for other thresholds). Clearly, rare and prevalent bacteria cluster out separately and constitute the largest part of the microbiome (in terms of number of species) but we can also notice a higher level of dependency between practically all regimes (see e.g. large connected component #1 for male and donorB). We present more examples (colored by abundance, occurrence, taxonomy, PC loading, seasonality, stationarity, and more) and discussion in the Supplement. In Fig. 5B-C we show the time evolution of bacteria in different components (cloud, largest connected components, the rest) stratified by the longitudinal regime for male and donorA subjects respectively (see also Supplementary Fig. 16A-B for other subjects). First, the cloud of singletons in each subject is dominated (in terms of total counts) by rare and temporal species (see Fig. 5C). Similarly, the largest connected component contains all regimes apart from rare and noise features that form a large cluster in the center (Supplementary Fig. 16A). However, the difference between the cloud and connected components is not obvious and is subject-dependent (threshold put on | *ρ* | also matters but doesn’t change qualitatively the results - see Supplementary Fig. 16C-D). Second, temporal and unstable species have a large effect on microbiome dynamics, again, irrespective on the dataset. In Fig. 5D we present the PC1+PC2 loading (the meaningfulness of a given region) of the cloud, the largest connected components and the rest as a function of *ρ*_thr_. Clearly, the cloud tends to be more important for explaining the microbiome variability with increasing *ρ*_thr_, but, even more importantly, the size of this effect is subject-independent.

## Discussion

In this study, we leveraged four long and dense time series of the gut microbiome in generally healthy individuals to elucidate its temporal dynamics. Our findings confirm subject-specific microbial signatures^29–32^. Through the assessment of fluctuations in alpha diversity over time, we provide evidence that the gut microbiome behaves in a non-stochastic manner as a unified entity, displaying stationarity and offering predictability based on its previous states. Additionally, our analyses reveal that at the taxonomic level, each individual’s gut microbiome is primarily composed of a few dominant groups of bacteria, with occasional temporal blooms of rare bacteria. Despite the small number of analyses of healthy gut microbiome behavior in time, our results are consistent with previous findings that the gut microbiome is host-unique and that its composition is stable over time^6, 8, 19, 33–35^.

Our study demonstrates the presence of an underlying seasonal pattern in the human gut microbiome. Through the application of the Fast Fourier Transform (FFT), we identify the existence of multiple dominant seasonalities within the gut microbiome. Moreover, we establish that by utilizing the seasonal component, it becomes possible to predict changes in the human gut microbiome over time with satisfactory performance. The gut microbiome is known to be strongly influenced by external factors, including diet^36^. Previous research on the seasonality of the gut microbiome has largely focused on specific datasets from isolated religious groups or indigenous hunter-gatherer communities, whose dietary habits are closely linked to seasonal changes in weather^37–40^. We hypothesize that the observed seasonal patterns in the gut microbiome may arise from intricate metabolic interactions among bacteria, which depend on various energy sources derived from the diet and/or their interactions with the external environment. The fluctuations in nutrient availability and dietary composition across seasons could potentially influence the growth and activity of specific bacterial groups, leading to the emergence of distinct seasonal behaviors. Further investigations into the metabolic pathways and interactions within the gut microbiome are warranted to elucidate the underlying mechanisms contributing to these observed seasonal patterns^18^. However, without pertinent metadata, annotations of specific bacterial functions (e.g. derived from shotgun metagenomics experiments), or more ubiquitous longitudinal study designs, discerning the precise origins of these seasonal fluctuations remain challenging.

Next, to define how particular bacteria behave in time, we described each bacteria with a longitudinal characteristics vector (Table 2). We analyze the abundance of bacteria exhibiting specific temporal behavior within the gut microbiome of each individual. We create a correlation matrix of longitudinal features to derive groups of features that exhibit similar behavior. Finally, we define 3 large groups of bacterial behavior in time: (1) the stable microbiome, (2) the volatile microbiome, and (3) noise. We show that the data from the human gut microbiome is, in terms of relative abundance, mostly noise that, we hypothesize, might derive from technical factors. Then, we show that in all four subjects, there exists a small fraction of bacteria that despite being few are highly abundant and are present in more than 90% of the time in the human gut microbiome. Finally, we show that there exists a group of volatile bacteria that, we hypothesize, react more vividly to environmental changes. Our results are cohesive with previous S. Gibbons research that the human gut microbiome consists of predictable autoregressive bacteria and stochastic non-autoregressive bacteria and that diet might be a key player in gut microbiome dynamics ^4,18^.

Graph analysis (performed with NetworkX) revealed that we can additionally group bacteria in a generally healthy human gut microbiome based on its co-occurrence relationships: noisy bacteria (the largest part in terms of a total number of species) that cluster out together, abundant bacteria (largest part in terms of a total number of counts) that presumably drive microbiome dynamics and a sizeable part of mostly singletons (denoted as “cloud”) that are not (anti)correlated with anything else. The last two regions are highly heterogeneous in terms of their longitudinal regime which may indicate complex relationships between bacteria from different regimes. The cloud seems to be the most intriguing, especially taxa classified as stable and prevalent that are present in it. However, higher quality data (larger taxonomic resolution) would be needed to analyze its dynamics and related functions.

Our study aligns closely with previous research, highlighting the coherence of our findings. What distinguishes our approach is the use of rigorous statistical methods, machine learning algorithms, econometric analysis, and graphical tools to examine the behavior of individual bacteria in the human gut. This allows other scientists to more efficiently quantify even large quantities of bacteria and gain new insights into the composition of the human gut microbiome community. We believe that our study facilitates scientists in understanding the behavior of bacteria in the human gut and aids in the development of predictive models. Traditionally, researchers have employed a methodology wherein they analyze the top 10% of the most frequently observed bacterial taxa to gain a comprehensive understanding of the microbiome dynamics over time^4,18,41^. However, our findings demonstrate that while this approach is indeed valuable and the removal of rare bacteria serves as an effective means of reducing dimensionality and noise, it is imperative to acknowledge the presence of bacteria that exhibit temporal patterns, emerging periodically in response to specific conditions within the host’s gastrointestinal tract.

Our research could also be valuable in designing gut microbiome studies and planning sampling strategies. The existence of these temporal bacteria strongly advocates for more frequent incorporation of temporal series analysis, as relying solely on bacteria sampled at a single time point may inadvertently overlook the presence of meaningful taxa. Such taxa, if disregarded, could potentially impact the accuracy of classification or regression models employed in various research applications.

Despite the valuable insights provided by our study, there are certain limitations that should be acknowledged. Firstly, in order to fully understand the dynamics of the microbiome, including seasonality, predictability, and change points, we require additional metadata beyond what was available for the studies we analyzed. This includes factors such as environmental conditions, dietary habits, or health status, and also collecting data following accepted metadata standards^42–44^. Secondly, to comprehensively annotate the functions of the microbiome, shotgun sequencing data is necessary. While our study focuses on ASVs, it is important to note that sequences and taxonomy alone may be less robust and informative compared to functions. This is because multiple taxa can potentially contribute to the same functions, and different individuals with distinct microbiomes may still exhibit functional similarities^35,45^.

Therefore, further investigations incorporating shotgun sequencing data and associated metadata are crucial for a more comprehensive understanding of the gut microbiome.

## Methods

### Data preparation

#### Datasets

For all analyses in this study we utilized two publicly available 16S sequencing gut microbiome datasets containing data of human gut microbiome from four, generally healthy adult individuals. Datasets were downloaded from the Qiita repository (https://qiita.ucsd.edu/). Demultiplexing, trimming, and feature table preparation were done using the Qiita framework. Missing time points were interpolated using PCHIP interpolation (see below for details). Interpolated data were rarefied to 18000 sequence count threshold. Rarefaction was performed using QIIME 2 (see below for details).

#### Dataset #1 Moving pictures of the human microbiome^7^

The first dataset utilized in this study comprises longitudinal 16S sequencing data obtained from the human microbiota. The dataset encompassed two individuals and covered four different body sites. The first individual was a generally healthy adult male who was sampled at four body sites for a duration of 443 days. The second individual, a generally healthy adult woman, was sampled for 185 days.

#### Dataset #2 Host lifestyle affects human microbiota on daily timescales^8^

The second dataset used in this study comprises longitudinal 16S sequencing measurements of the human gut and salivary microbiota dynamics for two generally healthy adult males. The data covers a duration of one year, with the first individual sampled for 365 days and the second individual sampled for 252 days.

#### Data preprocessing

For all datasets, raw data underwent preprocessing steps using the Qiita pipeline. These steps included demultiplexing, trimming the sequences to a standardized length of 100 nucleotides, and feature table preparation utilizing the Deblur algorithm. The resulting feature tables were then downloaded in Biom format, facilitating subsequent downstream analyses. For phylogenetic analysis, representative sequences were extracted from the dataset in FASTA format. Notably, only samples pertaining to the gut microbiome were included in this particular analysis, focusing on the microbial composition specifically within the gut.

#### Interpolation

In all four-time series, missing time points were interpolated using PCHIP (Piecewise Cubic Hermite Interpolation). PCHIP interpolation is a method that approximates a smooth curve or function between data points. It fits a cubic polynomial between adjacent points while ensuring the continuity of the function and its derivative. PCHIP is well-suited for microbiome data analysis as it maintains abundances above zero, preserves monotonicity, and avoids overshooting in cases of non-smooth data. Interpolation was performed using SciPy v.1.7.3 Python package with default settings.

#### Rarefaction

After interpolation, all four-time series underwent rarefaction to 180,000 sequences per sample to mitigate the influence of sequencing depth on alpha diversity analysis. Rarefaction was executed using the QIIME 2 v.2022.2.1 (https://docs.qiime2.org/2022.2.1/), a comprehensive software package designed for microbiome data analysis.

### Whole community analysis

#### PCoA between subjects

Aitchison distance between timepoints among individuals were calculated. Aitchinson distance was calculated on non-rarefied data after interpolation. The Aitchison distance is a statistical measure used to quantify compositional differences in relative abundance data, accounting for the constrained nature of compositional space^28^. The Aitchison distance is the Euclidean distance between compositions that have been transformed using the CLR (centered log-ratio) transformation. It possesses desirable properties such as scale invariance, perturbation invariance, permutation invariance, and sub-compositional dominance, which are not present in the standard Euclidean distance^46^. Aitchinson distance matrix was created from calculated distances between the time points of each individual. Next, Principal component Analysis (PCoA) was used on the distance matrix to reduce data dimensionality. PCoA is a dimensionality reduction technique used to visualize and explore patterns in multivariate data. It converts a distance or dissimilarity matrix into a set of orthogonal axes called principal coordinates, where each axis represents a linear combination of the original variables. Finally, using seaborn v.0.11.2 and Matplotlib v.3.1.3 Python packages we visualized the first two dimensions of the PCoA results to gain insights into the dissimilarities among the time series.

#### PCoA on individual subject

Aitchison distance between time points within the same individual was calculated. Aitchinson distance was calculated on non-rarefied data after interpolation. Next, PCoA was used to reduce the dimensionality of data, and the two first components were visualized using seaborn v.0.12.1 and Matplotlib v.3.5.3 Python packages.

#### Phylogenetic tree preparation

Phylogenetic tree construction was performed using QIIME 2 framework pipeline. The pipeline begins by using the mafft program to align representative sequences. Next, the pipeline filters the alignment to remove highly variable positions, which can introduce noise to the phylogenetic tree. Then, FastTree is used to generate the phylogenetic tree from the masked alignment. Finally, midpoint rooting is applied to position the tree’s root at the midpoint of the longest tip-to-tip distance in the unrooted tree. For this analysis, we use a rooted tree.

#### Alpha diversity Calculation

For each of the 4 individuals, we computed the Shannon diversity index and Faith’s diversity index on the rarefied gut microbiome data. Shannon Diversity index^47^ measures the richness and evenness of species in a community, focusing on species abundance distribution: 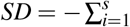 *p*_*i*_ log *p*_*i*_, where *s* is the number of amplicon sequence variants (ASVs) and *p*_*i*_ the proportion of the community represented by *i*th ASV. Faith’s Phylogenetic Index^48^ incorporates phylogenetic relatedness among species, emphasizing the evolutionary diversity of the community: *PD*_*i*_ = ∑ _*j∈ ℐ*_ *I*_*ij*_ branchlen _*j*_(*ℐ*), where *PD*_*i*_ is Faith’s phylogenetic diversity for sample *i, I*_*i j*_ indicates if sample *i* has any features that descend from node *j*, and branchlen _*j*_(*ℐ*) indicates the length of the branch to node *j* in the tree *ℐ*. Faith’s phylogenetic index was computed utilizing a rooted tree that was generated following the instructions outlined in the “Phylogenetic tree preparation” section. By constructing the rooted tree using the specified methodology, Faith’s phylogenetic index could be accurately calculated and applied to assess the phylogenetic diversity within the studied dataset. Alpha diversity indexes were calculated within QIIME 2 2022.2.1 framework.

#### Volatility

For each subject’s alpha diversity (Shannon diversity index and Faith’s phylogenetic index), the volatility was defined as the average conditional variance observed throughout the entire time series. The conditional variance was obtained by fitting a Generalized Auto-Regressive Conditional Heteroskedasticity GARCH(1, 1) model to the time series, with the model parameters determined through maximum likelihood estimation (MLE). The choice of 1, 1 for the GARCH model parameters was based on the observation that higher values did not yield improved results. The GARCH model was fitted using the arch v.5.3.1 Python package.

#### Trend analysis

Trend was calculated using a linear regression model, where time is the explanatory variable and alpha diversity index is the response variable. The explanatory variable was standardized such that it has 0 mean and variance of 1. We define the trend of the alpha diversity index as a regression coefficient of time. The linear regression model and data scaling were calculated using scikit-learn v.1.0.2 Python package.

#### Taxonomy analysis

Taxonomy was assigned to interpolated and rarefied data. To assign taxonomy we first trained a Naive Bayes classifier on the GreenGenes2 2022.10 database. Next, we assigned taxonomy to each subject’s sequences. Training of the classification as well as taxonomy assignment was performed within QIIME 2 framework. Plotting was performed using the seaborn v.0.11.2 python Package

#### Autocorrelation

For each subject the autocorrelation coefficient for 70 lags was calculated on alpha diversity variables (Shannon diversity index and Faith’s phylogenetic index). 95% confidence intervals were used to assess the statistical significance of the autocorrelation coefficient. The selection of 70 lags in gut microbiome data is somewhat arbitrary, as it was primarily chosen to effectively demonstrate the seasonal fluctuations. However, the choice of the specific number of lags can be subjective and dependent on individual preferences and research objectives.

#### Partial autocorrelation

For each subject partial autocorrelation coefficient for 70 lags was calculated on the alpha diversity variable (Shannon diversity index and Faith’s phylogenetic index). 95% confidence intervals were used to assess the statistical significance of the partial autocorrelation coefficient. Both autocorrelation and partial autocorrelation coefficients were calculated using statsmodels v.0.13.5 Python package with default settings and a significance level of 0.05.

#### Ljung-box test

For each subject the autocorrelation test was run on alpha diversity variables (Shannon diversity index and Faith’s phylogenetic index). The Ljung-Box test^49^ is a statistical test used to determine whether a time series is autocorrelated. The lag parameter was initially set to 30, offering a solid default value. Nevertheless, users have the freedom to customize it based on the length of their time series for optimal results. For each lag, we assume that the autocorrelation is present if the p-value is below the significance level of 0.05. The Ljung-Box test was performed using the statsmodels v.0.13.5 Python package with default settings and a significance level of 0.05.

#### Unit root tests

To analyze whether the microbiome is stationary as a whole we run two unit root tests^50^ - the KPSS (Kwiatkowski-Phillips-Schmidt-Shin) and ADF (Augmented Dickey-Fuller) on both alpha diversity measures (Shannon diversity index and Faith’s phylogenetic index). The KPSS test examines whether a time series exhibits trend or non-stationarity, while the ADF test determines whether a unit root is present, indicating non-stationarity. Both tests were performed using the statsmodels v.0.13.5 Python package with default settings and a significance level of 0.05.

#### Spectrum analysis

For each subject to detect repetitive patterns in alpha diversity (Shannon diversity index and Faith’s phylogenetic index) we used Fast Fourier Transform (FFT)^51^ to detect dominant seasonalities. FFT is an efficient algorithm used to transform a time-domain signal into its frequency-domain representation. It allows for the rapid computation of the discrete Fourier transform (DFT) by exploiting symmetries and redundancies in the data. To ensure result generalization, we focused on the initial 150 days of each individual’s time series. This duration was chosen because it represents the shortest length where no significant events, such as a period of diarrhea affecting the microbiome composition in donorB, occurred. We opted to remove this noisy period to enhance the accuracy of our analysis. First, we removed the trend from the data. To this purpose, linear regression model was fitted as described in “Trend” section. Then, the obtained trend was subtracted from the variable. Next, we ran FFT on detrended data. FFT results were plotted using a spectrogram, where on the x-axis we show the period in units of days, and on the y-axis we show period amplitude (PSD). All analyses for this part were done using SciPy v.1.7.3 Python package.

#### Flatness score

The spectrum flatness score measures the relative balance between the harmonic and non-harmonic components in a signal’s frequency spectrum, providing an indication of how “flat” or “noisy” the spectrum is. For each subject’s alpha diversity we calculated flatness score to asses its stochasticity. First, we detrended each time series by fitting a linear regression model to it and subtracting the predicted trend. Then, we calculated flatness score of the detrended time series using the spectrogram analysis. The flatness score was calculated using librosa v.0.10.0 Python package with default settings apart from the n_fft (FFT window size) parameter that was set to the half of the time series length.

#### FFT reconstruction

Alpha diversity seasonalities for each subject were sorted based on their amplitude. Starting from the seasonality with the highest amplitude, we employed the Inverse Fast Fourier Transform (IFFT) function to reconstruct the alpha diversity by considering only its *N* dominant seasonalities. Subsequently, we calculated the seasonal reconstructions score between the raw alpha diversity index and the alpha diversity index reconstructed using only its dominant seasonality. This analysis was performed for up to 10 dominant seasonalities. To visualize the relationship between the number of seasonalities used for signal reconstruction and the seasonal reconstructions score, we plotted the data using the Python seaborn package. IFFT was calculated with SciPy v.1.7.3 Python package. Spearman correlation coefficient was calculated with statsmodels v.0.13.5 Python package.

#### Seasonal reconstruction score

We define as seasonal reconstruction score Spearman correlation coefficient between raw and time series reconstructed using only its dominant seasonality.

#### Alpha diversity prediction

Dynamic Autoregressive Integrated Moving Average (ARIMA) model with a seasonal wave generated using Fast Fourier transform was used to forecast the behavior of alpha diversity over time. The training dataset consisted of the initial 80 days for each subject. The selection of ARIMA parameters (p, d, and q) and the determination of the number of seasonalities required to create the seasonal wave were performed using a grid search approach. To assess the performance of the models, we utilized the Mean Average Percentage Error (MAPE) and Wasserstein distance to measure the similarity between the predicted and true values of alpha diversity. The model with the best performance was chosen for predicting the test dataset. For each time series, we predicted the remaining time points and subsequently computed MAPE and Wasserstein distance between the true and predicted values of alpha diversity. To gain insights into the predictability of alpha diversity solely based on the alpha diversity index, we conducted cross-validation by considering consecutive intervals of 20 days. This enabled us to identify periods that were more challenging to predict. By calculating MAPE and Wasserstein distance in this manner, we obtained an evaluation of model performance on the test set. To fit ARIMA model statsmodels v.0.13.5 Python package was used. Wasserstein distance was calculated using SciPy v.1.7.3 Python package. Mean Average Percentage Error was calculated using scikit-learn v.1.0.2 Python package.

### Individual features

For each ASV we construct a longitudinal feature vector describing feature behavior in time. To investigate ASV we calculate its mean abundance, volatility, prevalence, loading, seasonality, trend, stationarity, and white noise behavior.

#### Mean abundance and standard deviation

For each ASV we define mean abundance as the mean number of reads per day in the whole time series, calculated on interpolated and rarefied data. Mean and standard deviation were calculated with NumPy v.1.21.6 Python package.

#### Prevalence

For each ASV we define mean prevalence as the percent of days where the number of reads for ASV is present compared to the whole time series length.

#### Loading

For each ASV we define mean loading as the loading derived from PCoA analysis of data for each subject (see PCoA on individual subject). For each subject, we first calculate Aitchinson’s distance between all time points. Next, we use PCoA to reduce the dimensionality of data into two components PC1 and PC2 respectively. Feature loading refers to the cumulative contribution or influence of individual variables (features) on two resulting coordinate axes.

#### Seasonality

To evaluate the stationarity of each ASV, we initially conducted unit root tests using the Kwiatkowski-Phillips-Schmidt-Shin (KPSS) and Augmented Dickey-Fuller (ADF) tests. These tests served to determine whether the ASVs exhibited characteristics of stationarity or non-stationarity. When KPSS test categorized the ASV as non-stationary, while the ADF test categorized it as stationary, the ASV underwent a detrending process to remove any underlying trend. Conversely, if the KPSS test deemed the ASV as stationary and the ADF test confirmed stationarity, the ASV was differenced by computing the differences between consecutive observations. In cases where both tests indicated non-stationarity, differencing was applied to the ASV. These procedures aimed to enhance the stationarity of the ASV for further analysis and modeling. For each stationary ASV we find dominant seasonalities using Fast Fourier Transform (FFT). Using 6 dominant seasonalities we use Inverse Fast Fourier Transform (IFFT) to create ASV fluctuation in time using only dominant seasonalities. Next, we computed seasonal reconstruction score between raw and seasonally reconstructed ASV trajectory. We define an ASV as seasonal if the correlation coefficient for maximally 6 seasonalities is above 0.4. We decided on this threshold based on seasonality analysis shown in Supplementary Fig. 10. FFT, IFFT, and correlation coefficients were run using SciPy v.1.7.3 Python package with default settings.

#### Trend

For each ASV trend was calculated using a linear regression model, where time is the explanatory variable and ASV fluctuation in time is the response variable. The explanatory variable was standardized that it has 0 mean and variance of 1. We define the trend of the ASV as a regression coefficient of time. The linear regression model data scaling were calculaled using scikit-learn v.1.0.2 Python package.

#### Stationarity

To define if a given ASV is stationary we used two unit root tests (KPSS and ADF – see previous subsection for details). A time series is defined as stationary if both tests confirm that it does not contain a unit root. Both tests were performed using the statsmodels v.0.13.5 Python package with default settings and a significance level of 0.05.

#### White noise behavior

Our subjective definition of white noise behavior is based on two criteria: the absence of autocorrelation and a high flatness score. To assess these criteria, we performed two statistical analyses: 1) the Ljung-Box test statistics was computed for each ASV’s 30 lags using the SciPy v.1.7.3 Python package with the default settings and significance level of 0.05; 2) the flatness score was calculated for each ASV separately using the librosa v.0.10.0 Python package, also using default settings. To establish a threshold for the flatness score indicating random behavior, we plotted the Ljung-Box test p-values against the flatness score. We determined a flatness score threshold of 0.4. Thus, we define a time series as demonstrating white noise behavior if the absence of autocorrelation is validated by the Ljung-Box test, with a p-value *>* 0.05 for all lags and its flatness score exceeds 0.4. For threshold analysis see Supplementary Fig 6.

#### Correlation matrix

We created a Spearman correlation coefficient matrix between longitudinal features using combined data for all ASVs from all four subjects. The Spearman correlation coefficient was calculated using SciPy v.1.7.3 Python package.

### Cluster analysis

Graphs in Fig. 5 (and in Supplementary Fig. 15) have been generated using the NetworkX v.2.8.4 Python package (https://networkx.org/) using spring layout (`spring_layout` method) with default parameters. Input matrix *D* has been prepared as follows: 1) for a given dataset raw counts (after interpolation; for donorB only first 150 days have been taken into account) were transformed using centered log-ratio transformation with pseudocunt equal to 1 using `skbio.stats.composition.clr` method (scikit-bio v.0.5.6); 2) all-vs-all proportionality matrix *ρ* has been computed as (NumPy v.1.23.5): `rho(x, y) = 1 - numpy.var(x - y) / (numpy.var(x) + numpy.var(y));` 3) for a given *ρ*_thr_ (a parameter that controls which species co-occur on anti co-occur), the |*ρ*| matrix has been truncated i.e. all entries ≤ *ρ*_thr_ were zeroed; 4) finally, pseudo-distance matrix was defined as *D* = 1 −|*ρ*|.

In the paper we use *ρ*_thr_ = 0.6 which provides good separability between different subgraphs (compare with Supplementary Fig. 15L–M, corresponding to *ρ*_thr_ = 0.5 and 0.7 where relations between bacteria are more blurred). Certain quantitative results depend on *ρ*_thr_ but most of the qualitative conclusions are independent of that parameter (see discussion in the main text for details).

Hierarchical clustering discussed in the Supplement (see Supplementary Fig. 20 and 21) has been performed using `scipy.cluster.hierarchy` Python package (SciPy v.1.10.0). First, linkages were constructed using the linkage method with method=‘complete’ (other values of that argument have been also tested but didn’t perform that well). Second, clusters were created by cutting the linkage trees using cut_tree method with hc_height = 2 (again, higher values performed poorly).

## Acknowledgements

We would like to thank Dr. Ewa Szczurek and Marcin Moz?ejko from Warsaw University, Poland for useful discussions on the methodological framework of this work.

## Author information

### Contributions

T.K. and Z.K. conceived the study; Z.K. and P.S. conducted the experiments, Z.K., P.S. and T.K. analysed the results. All authors reviewed the manuscript. All authors read and approved the final manuscript.

## Ethics declarations

### Ethics approval and consent to participate

Not applicable.

### Consent for publication

Not applicable.

## Data and code availability

Code for reproducing figures in this manuscript and many useful methods’ implementations used in this work may be found here: https://github.com/bioinf-mcb/dynamo.

## References

1. Manor, O. et al. Health and disease markers correlate with gut microbiome composition across thousands of people. Nat. Commun. 11, 5206 (2020).

2. Shreiner, A. B., Kao, J. Y. & Young, V. B. The gut microbiome in health and in disease. Curr Opin Gastroenterol 31, 69–75 (2015).

3. Fan, Y. & Pedersen, O. Gut microbiota in human metabolic health and disease. Nat. Rev. Microbiol. 19, 55–71 (2021).

4. Shenhav, L. et al. Modeling the temporal dynamics of the gut microbial community in adults and infants. PLOS Comput. Biol. 15, e1006960 (2019).

5. Stein, R. R. et al. Ecological modeling from Time-Series inference: Insight into dynamics and stability of intestinal microbiota. PLOS Comput. Biol. 9, e1003388 (2013).

6. Rajilić-Stojanović, M., Heilig, H. G. H. J., Tims, S., Zoetendal, E. G. & de Vos, W. M. Long-term monitoring of the human intestinal microbiota composition. Environ Microbiol (2012).

7. Caporaso, J. G. et al. Moving pictures of the human microbiome. Genome Biol 12, R50 (2011).

8. David, L. A. et al. Host lifestyle affects human microbiota on daily timescales. Genome Biol 15, R89 (2014).

9. Poyet, M. et al. A library of human gut bacterial isolates paired with longitudinal multiomics data enables mechanistic microbiome research. Nat Med 25, 1442–1452 (2019).

10. Äijö, T., Müller, C. L. & Bonneau, R. Temporal probabilistic modeling of bacterial compositions derived from 16S rRNA sequencing. Bioinformatics 34, 372–380 (2018).

11. Ang, Q. Y. et al. Ketogenic diets alter the gut microbiome resulting in decreased intestinal th17 cells. Cell 181, 1263–1275.e16 (2020).

12. Delannoy-Bruno, O. et al. Evaluating microbiome-directed fibre snacks in gnotobiotic mice and humans. Nature 595, 91–95 (2021).

13. Roager, H. M. et al. Whole grain-rich diet reduces body weight and systemic low-grade inflammation without inducing major changes of the gut microbiome: a randomised cross-over trial. Gut 68, 83–93 (2017).

14. von Schwartzenberg, R. J. et al. Caloric restriction disrupts the microbiota and colonization resistance. Nature 595, 272–277 (2021).

15. Ianiro, G. et al. Variability of strain engraftment and predictability of microbiome composition after fecal microbiota transplantation across different diseases. Nat Med 28, 1913–1923 (2022).

16. Schmidt, T. S. B. et al. Drivers and determinants of strain dynamics following fecal microbiota transplantation. Nat. Medicine 28, 1902–1912 (2022).

17. Gupta, V. K. et al. A predictive index for health status using species-level gut microbiome profiling. Nat Commun 11, 4635 (2020).

18. Gibbons, S. M., Kearney, S. M., Smillie, C. S. & Alm, E. J. Two dynamic regimes in the human gut microbiome. PLoS Comput. Biol 13, e1005364 (2017).

19. Faith, J. J. et al. The long-term stability of the human gut microbiota. Science 341, 1237439 (2013).

20. Flint, H. J., Duncan, S. H., Scott, K. P. & Louis, P. Links between diet, gut microbiota composition and gut metabolism. Proc Nutr Soc 74, 13–22 (2014).

21. Peterson, C. T. et al. Short-Chain fatty acids modulate healthy gut microbiota composition and functional potential. Curr Microbiol 79, 128 (2022).

22. Yang, J. et al. Oscillospira - a candidate for the next-generation probiotics. Gut Microbes 13, 1987783 (2021).

23. Abdugheni, R. et al. Metabolite profiling of human-originated lachnospiraceae at the strain level. iMeta 1, e58 (2022).

24. Schupack, D. A., Mars, R. A. T., Voelker, D. H., Abeykoon, J. P. & Kashyap, P. C. The promise of the gut microbiome as part of individualized treatment strategies. Nat Rev Gastroenterol Hepatol 19, 7–25 (2021).

25. Bender, J. M. et al. Quantification of variation and the impact of biomass in targeted 16S rRNA gene sequencing studies. Microbiome 6, 155 (2018).

26. Hildebrand, F. et al. Dispersal strategies shape persistence and evolution of human gut bacteria. Cell Host Microbe 29, 1167–1176.e9 (2021).

27. Turnbaugh, P. J. et al. A core gut microbiome in obese and lean twins. Nature 457, 480–484 (2008).

28. Gloor, G. B., Macklaim, J. M., Pawlowsky-Glahn, V. & Egozcue, J. J. Microbiome datasets are compositional: And this is not optional. Front Microbiol 8, 2224 (2017).

29. Faust, K. et al. Signatures of ecological processes in microbial community time series. Microbiome 6, 120 (2018).

30. Alshawaqfeh, M., Serpedin, E. & Younes, A. B. Inferring microbial interaction networks from metagenomic data using SgLV-EKF algorithm. BMC Genomics 18, 228 (2017).

31. Pimm, S. L. & Redfearn, A. The variability of population densities. Nature 334, 613–614 (1988).

32. Fisher, C. K. & Mehta, P. Identifying keystone species in the human gut microbiome from metagenomic timeseries using sparse linear regression. PLoS One 9, e102451 (2014).

33. Costello, E. K. et al. Bacterial community variation in human body habitats across space and time. Science 326, 1694–1697 (2009).

34. Voigt, A. Y. et al. Temporal and technical variability of human gut metagenomes. Genome Biol 16, 73 (2015).

35. Human Microbiome Project Consortium. Structure, function and diversity of the healthy human microbiome. Nature 486, 207–214 (2012).

36. Gentile, C. L. & Weir, T. L. The gut microbiota at the intersection of diet and human health. Science 362, 776–780 (2018).

37. Davenport, E. R. et al. Seasonal variation in human gut microbiome composition. PLoS One 9, e90731 (2014).

38. Davenport, E. R. et al. Genome-Wide association studies of the human gut microbiota. PLoS One 10, e0140301 (2015).

39. Koliada, A. et al. Seasonal variation in gut microbiota composition: cross-sectional evidence from ukrainian population. BMC Microbiol 20, 100 (2020).

40. Sung, J. et al. Global metabolic interaction network of the human gut microbiota for context-specific community-scale analysis. Nat Commun 8, 15393 (2017).

41. Xiao, J. et al. Predictive modeling of microbiome data using a Phylogeny-Regularized generalized linear mixed model. Front Microbiol 9, 1391 (2018).

42. Vangay, P. et al. Microbiome metadata standards: Report of the national microbiome data collaborative’s workshop and Follow-On activities. mSystems 6 (2021).

43. Knight, R. et al. Best practices for analysing microbiomes. Nat Rev Microbiol 16, 410–422 (2018).

44. Navas-Molina, J. A., Hyde, E. R., Sanders, J. & Knight, R. The microbiome and big data. Curr Opin Syst Biol 4, 92–96 (2017).

45. Lloyd-Price, J. et al. Strains, functions and dynamics in the expanded human microbiome project. Nature 550, 61–66 (2017).

46. Quinn, T. P., Erb, I., Richardson, M. F. & Crowley, T. M. Understanding sequencing data as compositions: an outlook and review. Bioinformatics 34, 2870–2878 (2018).

47. Wagner, B. D. et al. On the use of diversity measures in longitudinal sequencing studies of microbial communities. Front Microbiol 9, 1037 (2018).

48. Armstrong, G. et al. Efficient computation of faith’s phylogenetic diversity with applications in characterizing microbiomes. Genome Res 31, 2131–2137 (2021).

49. Burns, P. Robustness of the Ljung-Box test and its rank equivalent. SSRN Electron. J. (2002).

50. Herranz, E. Unit root tests. WIREs Comput. Stat. 9, e1396 (2017).

51. Brigham, E. O. The Fast Fourier Transform and Its Applications (Prentice-Hall, Inc., USA, 1988).

