## Supplementary materials for "Microbiome time series data reveal predictable patterns of change"

#### Supplementary information

Zuzanna Karwowska[1,2], Pawel Szczerbiak[1], Tomasz Kosciolek[1]

1 Malopolska Centre of Biotechnology, Jagiellonian University, Krakow, Poland
2 Doctoral School of Exact and Natural Sciences, Jagiellonian University, Krakow, Poland

#### Table of contents

Whole community analysis 2
Individual features analysis 8
Cluster analysis 13
NetworkX graphs 13
Regime evolution in time 20
Examples 23
Sanity check - Euclidean distance vs |$\rho$| 24
Hierarchical clustering 25

### Whole community analysis

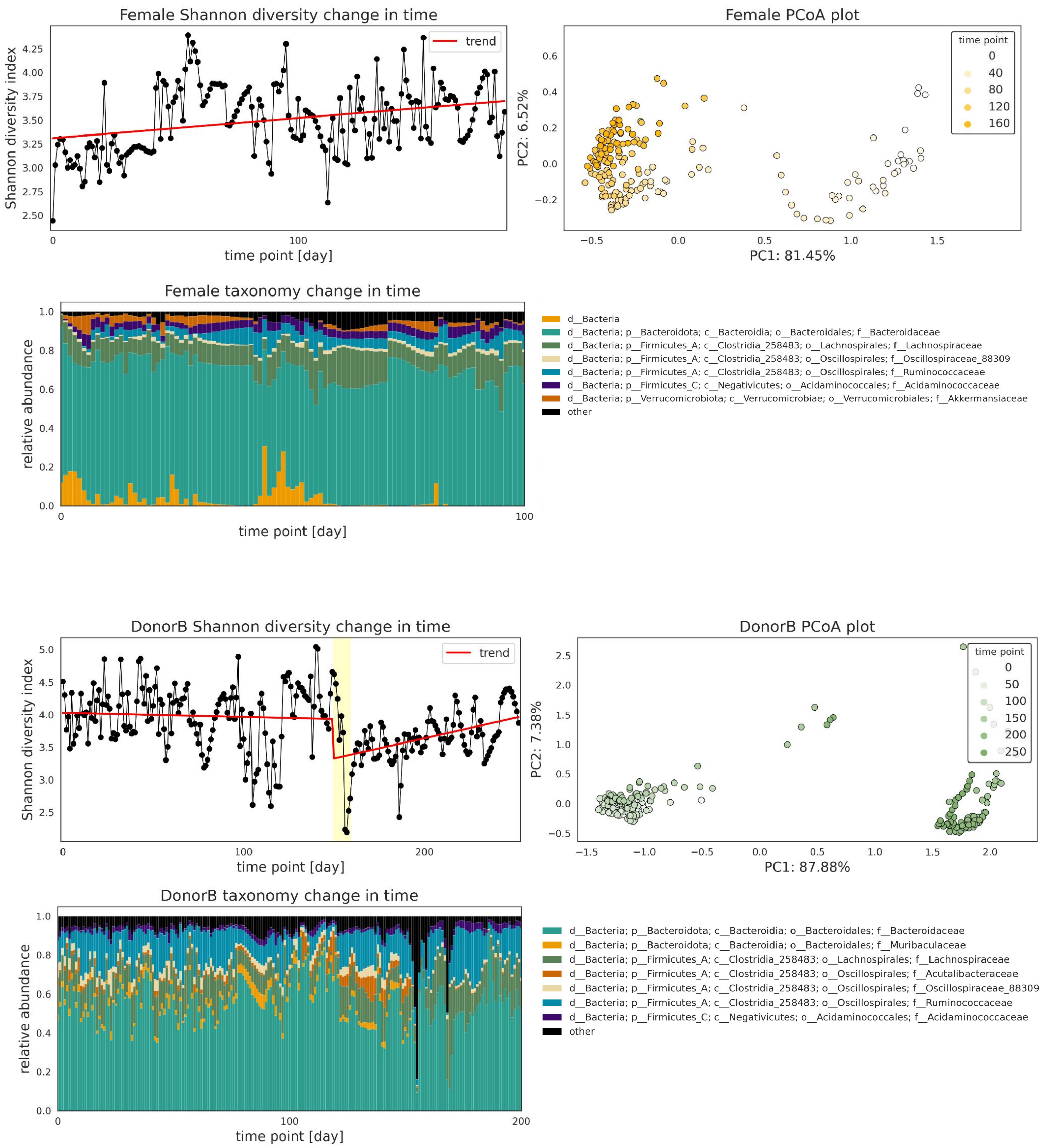


**Supplementary Figure 1. Overview of microbiome behavior in time** for donorB **(A)** and the female subject **(B)**. The female subject trend is growing, however, it can be seen from the PCoA analysis that the first 50 days are different from the rest of the time series and that the trend starting from day 50 is stable. From donorB metadata we know that he was suffering from diarrhea on days around day 150. We see that this event decreases his alpha diversity which drops around day 150 and then increases to the baseline level. Also during the event, we can see changes in taxonomy as new taxa appear.

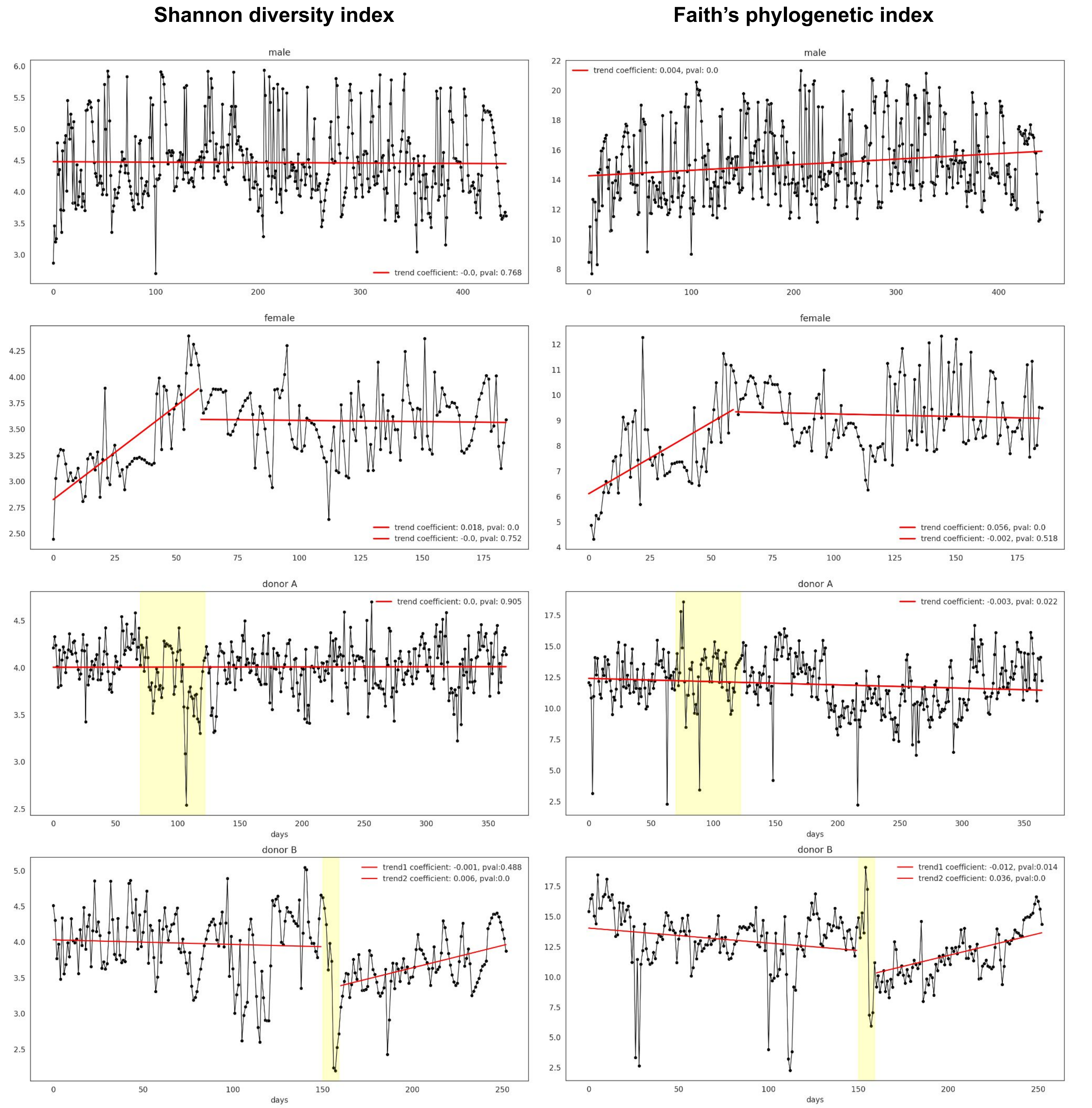


**supplementary Figure 2. Shannon diversity index and Faith's phylogenetic index behavior in time** (each row represents one subject). Red lines represent trend and trend coefficient that was calculated using regression analysis. In female subject we show that the trend becomes stable from day 60. We also show that donorA alpha diversity is stable in time despite perturbation.

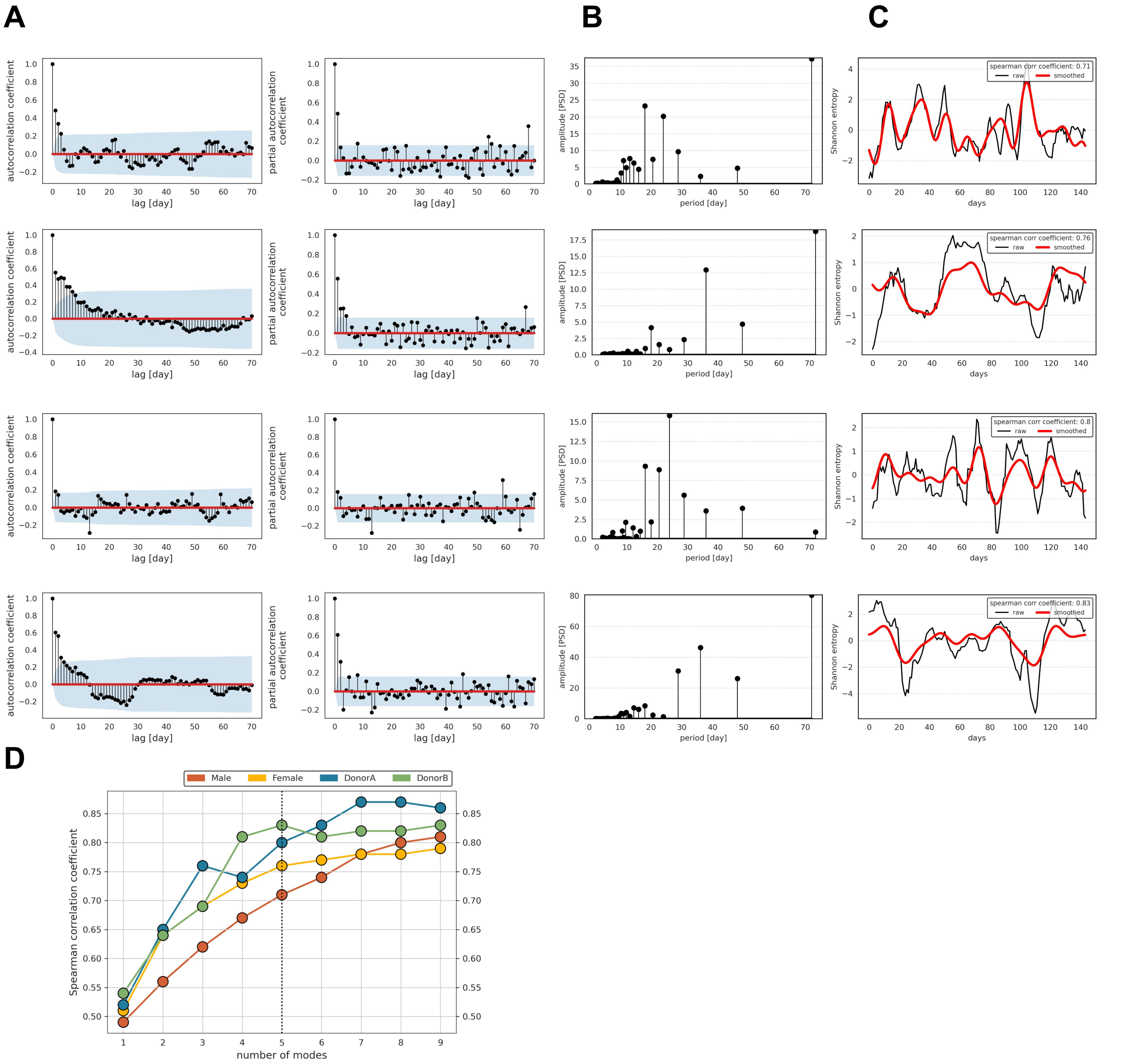


**Supplementary Figure 3. Faith's phylogenetic index behavior in time. A.** Autocorrelation coefficients plots (blue area shows the significance level). **B.** Spectrogram showing most dominant seasonalities of human gut microbiome. **C.** Reconstruction of alpha diversity using 5 dominant seasonalities plotted against raw alpha diversity change in time. **D.** plot showing the relationship between number of used seasonalities to reconstruct alpha diversity and the seasonal reconstruction score.

| Subject | Alpha diversity | Test | Test result |
|---|---|---|---|
| male | Shannon diversity index | ADF p value | 0.0 |
| | | KPSS p value | 0.1 |
| | | flatness score | 0.04 |
| | | Ljung-Box test | < 0.05 |
| female | | ADF p value | 0.008 |
| | | KPSS p value | 0.1 |
| | | flatness score | 0.01 |
| | | Ljung-Box test | < 0.05 |
| donorA | | ADF p value | 0.0 |
| | | KPSS p value | 0.1 |
| | | flatness score | 0.04 |
| | | Ljung-Box test | < 0.05 |
| male | Faith's phylogenetic index | ADF p value | 0.0 |
| | | KPSS p value | 0.1 |
| | | flatness score | 0.04 |
| | | Ljung-Box test | < 0.05 |
| female | | ADF p value | 0.048 |
| | | KPSS p value | 0.08 |
| | | flatness score | 0.02 |
| | | Ljung-Box test | < 0.05 |
| donorA | | ADF p value | 0.0 |
| | | KPSS p value | 0.1 |
| | | flatness score | 0.14 |
| | | Ljung-Box test | < 0.05 |
| donorB | | ADF p value | 0.0 |
| | | KPSS p value | 0.1 |
| | | flatness score | 0.0 |
| | | Ljung-Box test | < 0.05 |

**Supplementary Table 1. Statistical tests showing alpha diversity behavior in time:** unit root presence (ADF and KPSS tests), presence of autocorrelation (Ljung-Box test) and spectral flatness score. Both, Shannon diversity index and Faith's PD index do not have unit root, do have autocorrelation and have low flatness score which suggest that they behave in a non-stochastic manner.

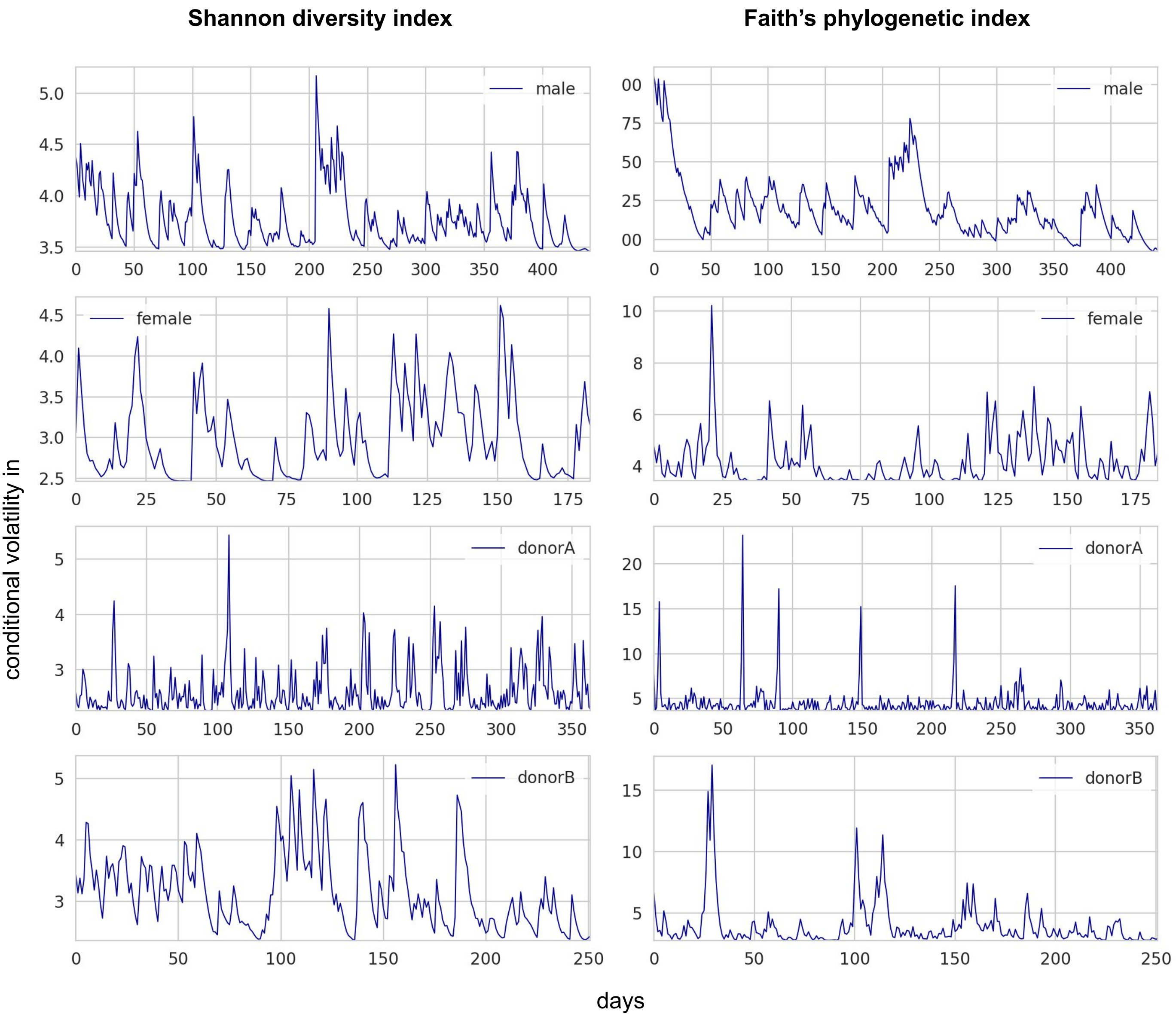


**Supplementary Figure 4. Line plots of conditional variance estimated using GARCH model** showing higher variance periods in each alpha diversity time series. Left: Shannon diversity index volatility; right: Faith's PD index volatility.

**A**

| Subject | p | d | q |
|---|---|---|---|
| male | 3 | 0 | 2 |
| female | 12 | 0 | 1 |
| donorA | 3 | 0 | 1 |
| donorB | 3 | 0 | 1 |

**B**

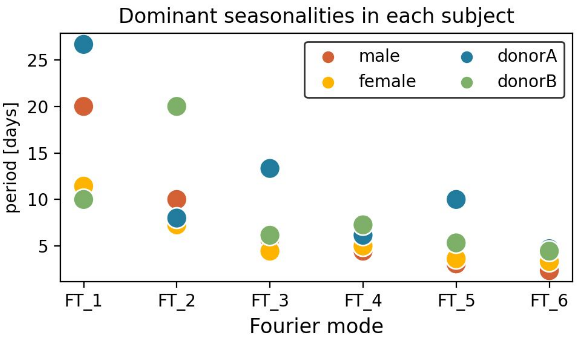


**C**

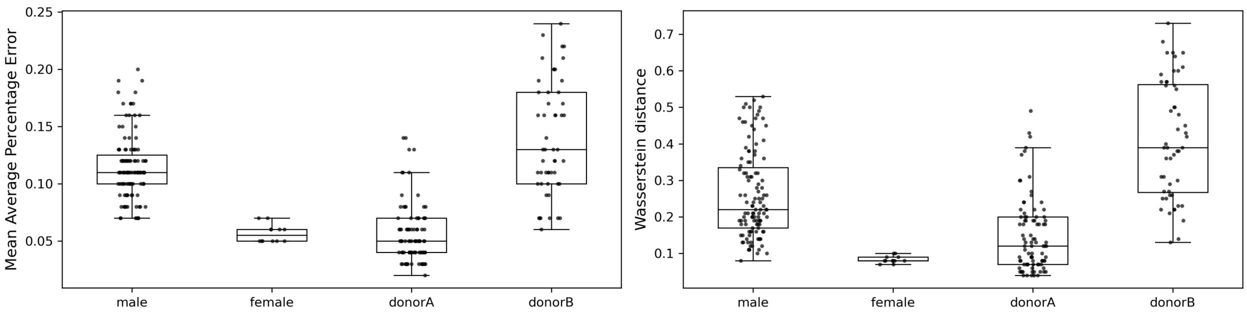


**Supplementary Figure 5. Dynamic Arimax model results. A.** ARIMA model parameters; **B.** Dominant seasonalities plot. Figure depicts the relationship between the number of seasonalities and their corresponding period lengths in constructing exogenous variables for the predictive model. The analysis includes multiple subjects, with a focus on donorA. The first Fourier mode for donorA has a 25-day period. The plot shows that the first mode has the longest period, and subsequent modes have decreasing periods with amplitude. Optimal seasonalities for male, female, donorA, and donorB were found to be 5, 3, 6, and 3, respectively. These findings emphasize the importance of considering seasonalities and their lengths when constructing exogenous variables for accurate predictions across subjects; **C.** The test set cross-validation results of the predictive model performance are presented, with the error measure calculated on a 20-day interval every 5 days. The first set of boxplots displays the mean average percentage error for each fold, while the second set demonstrates the Wasserstein distance. The analysis reveals that the model's performance is not consistently favorable across all folds. There are instances where the model does not perform well, indicating a lack of generalizability in its predictions. This lack of generalizability is attributed to external changes, such as events occurring in donorA and donorB. These findings suggest that the predictive model may be influenced by unpredictable factors or events specific to individual donors, making it challenging to achieve consistent and accurate predictions. Further investigation and refinement of the model may be necessary to enhance its generalizability and robustness to external variations.

### Individual features analysis

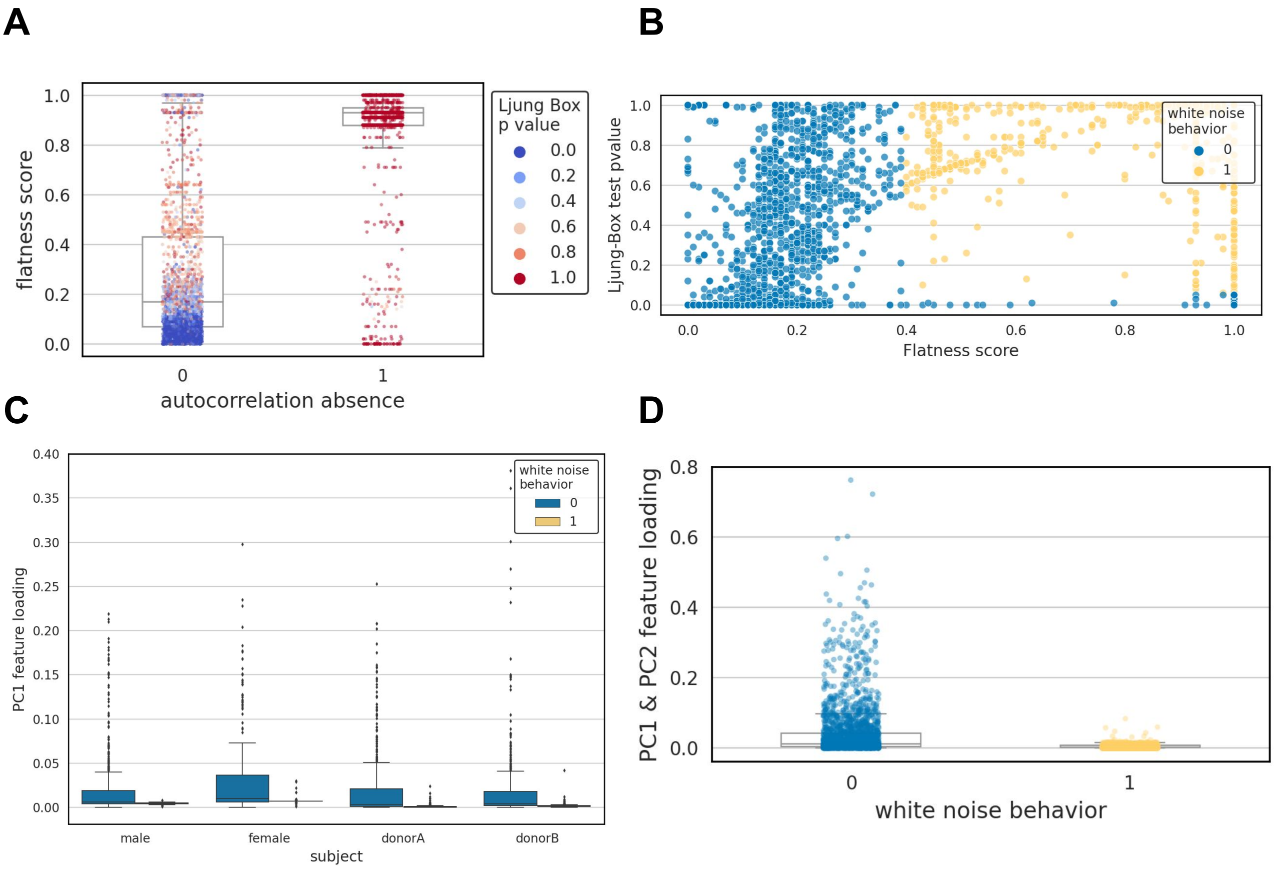


**Supplementary Figure 6. White Noise Definition. A.** and **B.** display the distribution of Ljung-Box test p-values against the flatness score for each taxon analyzed. These figures serve to illustrate the relationship between these two measures. Based on the observed distribution, we have chosen to introduce a new artificial measure termed "white noise behavior." This measure encompasses taxons that meet the following criteria: a Ljung-Box p-value less than 0.5 and a flatness score above 0.4. However, we recommend conducting a thorough analysis of such a distribution on individual datasets to determine the optimal threshold for defining white noise behavior. **C.** and **D.** provide additional insight into the impact of features identified as noise on data variance. This figure demonstrates that these identified features have minimal or no effect on the overall variance of the data.

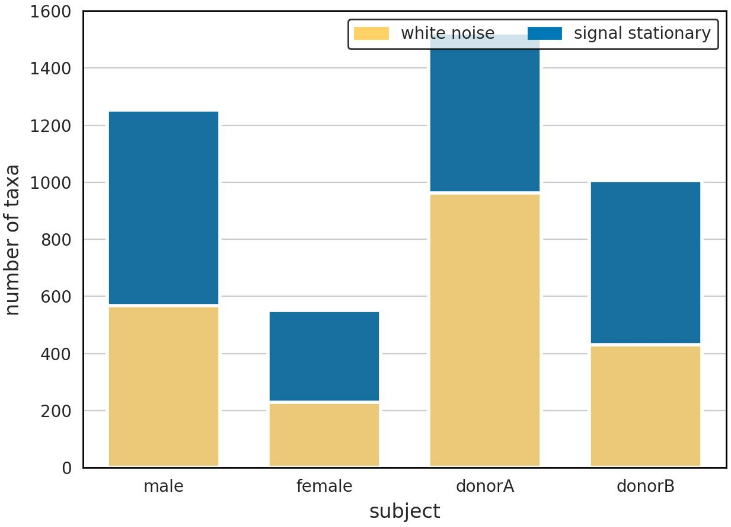


**Supplementary Figure 7.** Number of White noise bacteria in human gut microbiome.

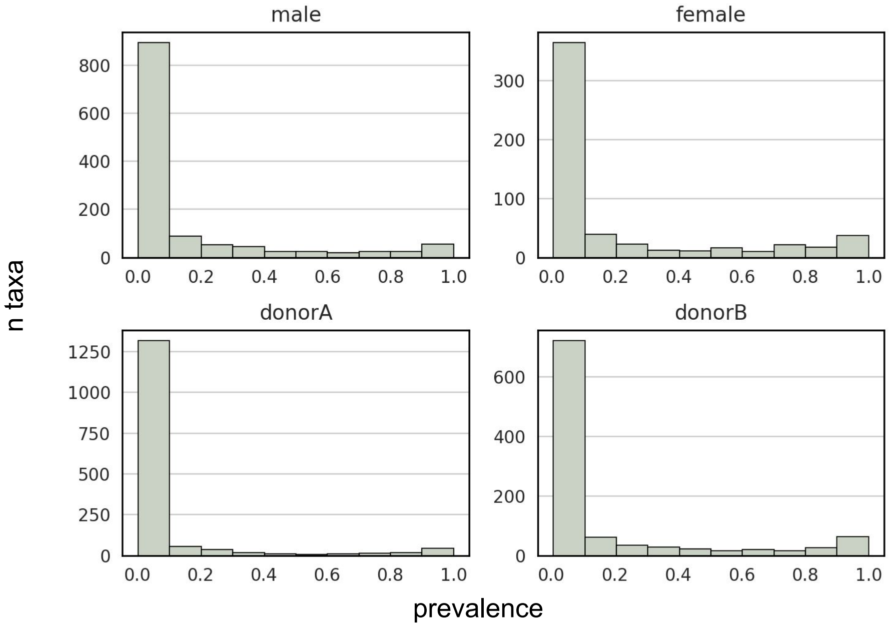


**Supplementary Figure 8. Taxa prevalence between subjects.** A significant proportion of bacteria were classified as rare or predominantly absent (they are present in less than 10% of time series), while a smaller fraction was consistently present in the gut microbiome. Another subset appeared intermittently, making them neither rare nor constantly present. Surprisingly, almost half of each individual's microbiome was classified as noise, likely resulting from technical factors and metabolic conditions.

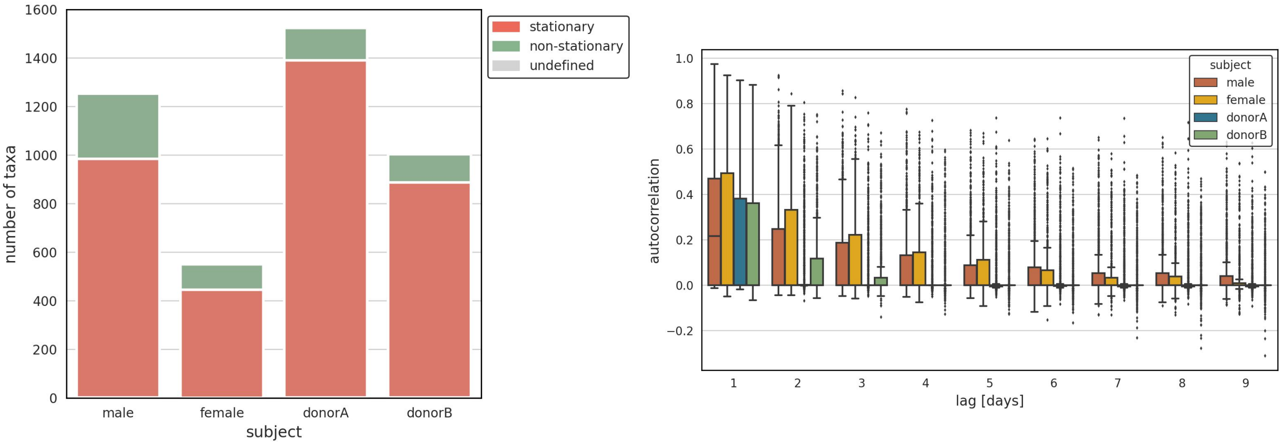


**Supplementary Figure 9.** Taxa stationarity and autocorrelation.

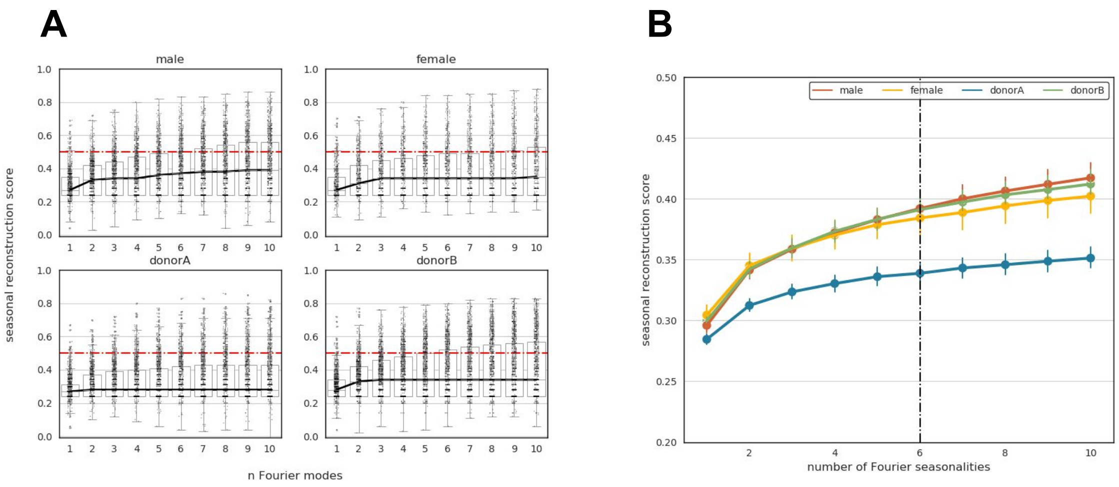


**Supplementary Figure 10. Number of seasonalities present in human gut microbiome. A.** The analysis of the required number of seasonalities needed to accurately reconstruct the raw bacterial fluctuation in time. Each data point represents a specific bacterial species. The x-axis represents the bacterial species, while the y-axis denotes the number of seasonalities required for reconstruction. **B.** Mean relationship between the number of modes used for reconstructing raw bacteria counts and the corresponding reconstruction score. The x-axis represents the number of modes, whereas the y-axis denotes the mean seasonal reconstruction score. The results demonstrate that, in contrast to alpha diversity, different bacterial species exhibit diverse behaviors. Additionally, the analysis highlights that the number of Fourier modes needed to accurately reconstruct the raw signal varies significantly among bacterial species (Panel A). These findings shed light on the complexity and heterogeneity of the gut microbiome seasonal dynamics.

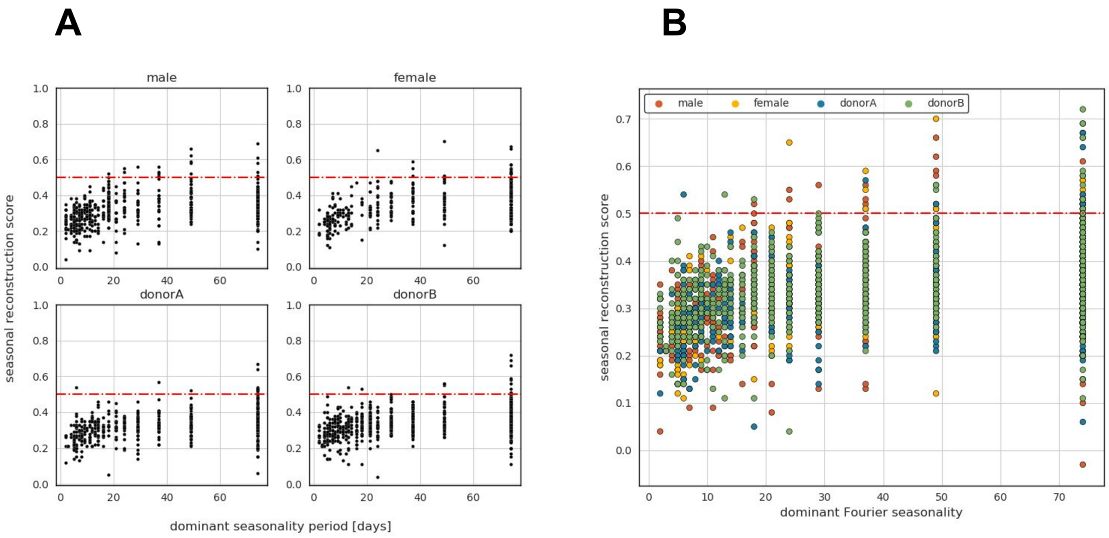


**Supplementary Figure 11. Dominant seasonalities in human gut microbiome. A.** Dominant bacterial seasonality and corresponding seasonal reconstruction score. The red line represents a score cutoff of 0.5. It is observed that several bacteria exhibit short seasonalities identified by FFT with low reconstruction scores, suggesting that they may be noise. Moreover, longer seasonalities tend to have higher reconstruction scores. **B**. The panel demonstrates that the seasonalities in the gut microbiome are generally similar across analysed subjects, indicating consistency in the observed patterns. Overall, the plot provides insights into the dominant seasonalities present in the human gut microbiome. The findings suggest the presence of meaningful seasonal patterns in certain bacteria, while cautioning against short seasonalities with low reconstruction scores. Additionally, the consistent patterns observed across subjects contribute to our understanding of the general characteristics of gut microbiome seasonal dynamics.

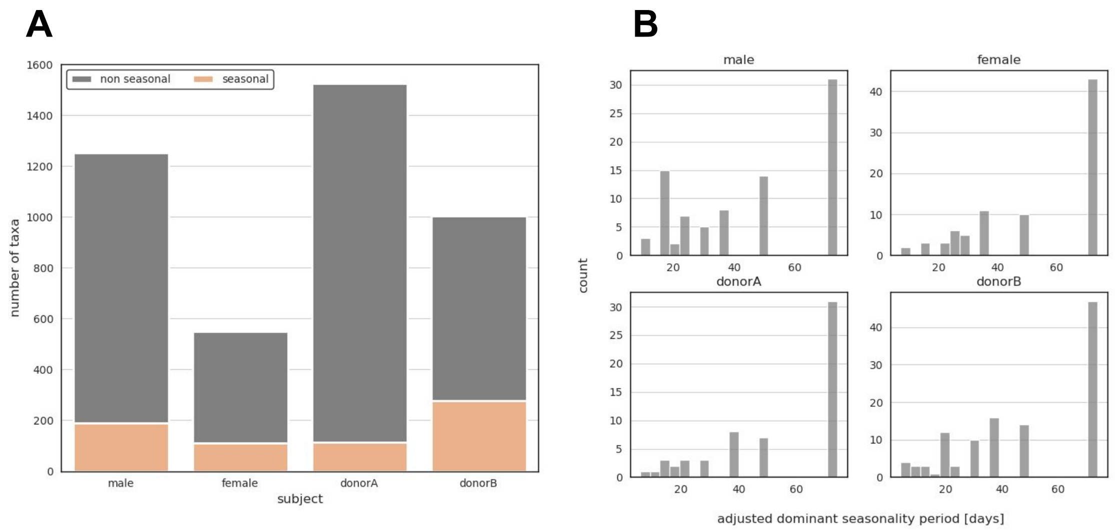


**Supplementary Figure 12. Number of seasonal bacteria in human gut microbiome. A.** Number of bacteria classified as seasonal in each subject. The classification is based on the successful reconstruction of bacterial raw counts using 6 Fourier seasonalities and a minimum seasonal reconstruction score of 0.5. This panel provides an overview of the prevalence of seasonal bacteria across subjects. **B.** Adjusted dominant seasonalities in bacteria for each subject. The adjusted seasonalities were identified based on a reconstruction correlation score of at least 0.4. This panel highlights the presence of more reliable and significant seasonal patterns in the bacterial population of each subject.

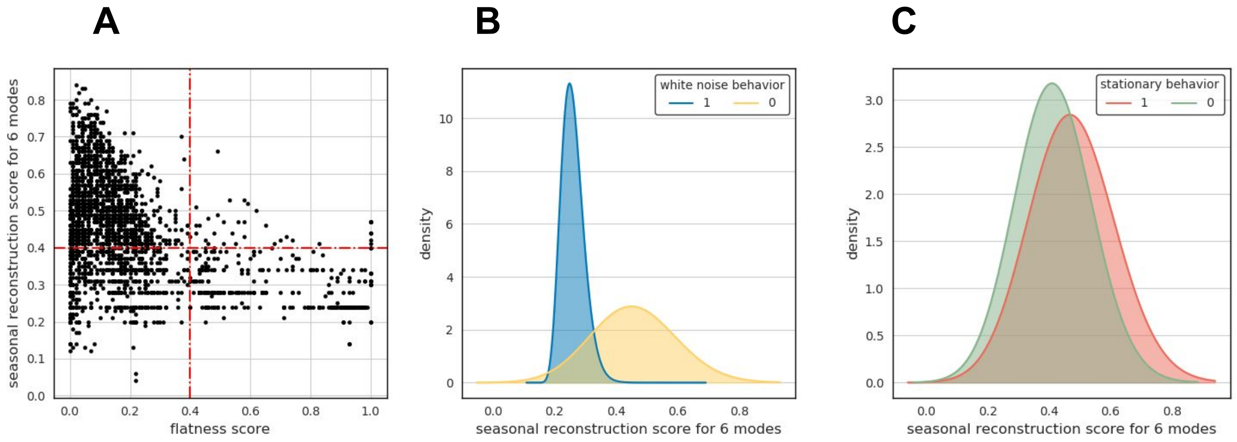


**Supplementary Figure 13. Different Characteristics of Seasonal Bacteria in the Human Gut Microbiome. A.** Relationship between the Seasonal Reconstruction Score of 6 Fourier Modes and Bacteria Flatness Score: This figure depicts the association between the seasonal reconstruction score, derived from the analysis of six dominant Fourier modes, and the bacteria flatness score. The results reveal that seasonal bacteria exhibit a remarkably low flatness score, indicating a high degree of regularity in their seasonal patterns. This observation serves as a sanity check, validating the presence of distinct seasonal behavior in these bacteria. **B.** Seasonal Reconstruction Score of 6 Fourier Modes for Bacteria Classified as White Noise: This panel showcases the seasonal reconstruction scores for individual bacteria, differentiating those classified as white noise from bacteria classified as 'signal'. Notably, bacteria displaying stochastic behavior, characterized by white noise patterns, generally do not exhibit discernible seasonality. Despite considering their six dominant seasonalities, we are unable to accurately reconstruct the seasonal patterns of these bacteria. This finding underscores the absence of predictable seasonal behavior in bacteria with stochastic dynamics. **C.** Relationship between Stationary and Seasonal Behavior: In this segment, we explore the connection between stationary and seasonal behavior in bacteria. The figure demonstrates that, on average, seasonal bacteria exhibit a stationary behavior, indicating a consistent and stable seasonal component over time. This observation suggests that the present seasonal component within these bacteria remains relatively unchanged and can be considered a stable feature.

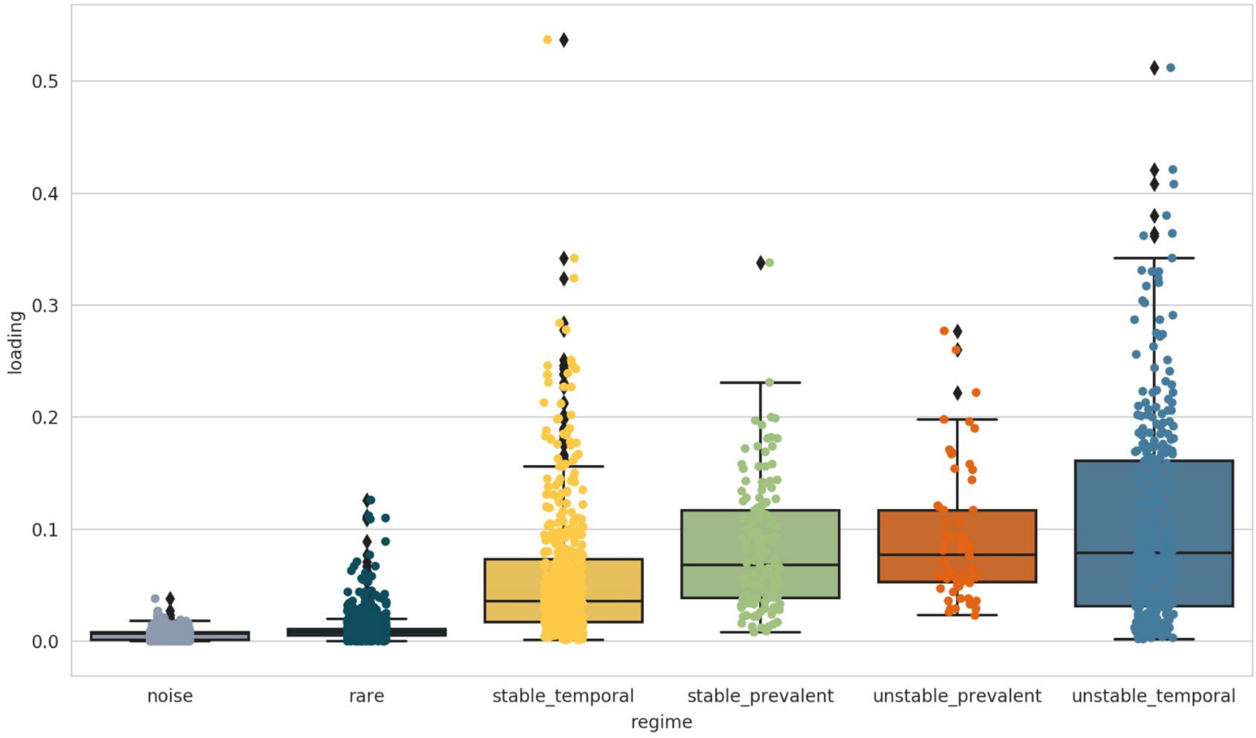


**Supplementary Figure 14. Cumulative PCoA feature loading of each bacterial regime.** Each point represents a bacteria displaying different regime. We can see that the bacteria belonging to rare and noise regime has the lowest regime despite being the most numerous. On the other hand unstable and temporal regime has the highest loading. PCoA was calculated for each subject separately based on it's Aitchison distance matrix between timepoints.

| Subject | All | Rarefied | Signal | Noise | Rare |
|---|---|---|---|---|---|
| male | 1399 | 1253 (90%) | 358 (26%) | 426 (30%) | 615 (44%) |
| female | 591 | 551 (93%) | 187 (32%) | 189 (32%) | 215 (37%) |
| donorA | 2862 | 1524 (53%) | 210 (7%) | 643 (23%) | 2009 (70%) |
| donorB | 1524 | 1005 (66%) | 280 (18%) | 318 (21%) | 926 (61%) |

**Supplementary Table 2.** Number of bacteria in each subject. All percentages are computed with respect to "All" (second column)l. Signal bacteria (stable / unstable and prevalent / temporal - see Supplementary Figure 14) comprises ~30% of male and female microbiomes and 7 / 18% of donorA / donorB microbiomes respectively (this may be an artifact coming from experiment). However, the absolute number is stable for all subjects and ranges between 187 and 358.

### Cluster analysis

#### NetworkX graphs

Panels in Supplementary Figure 15 (see caption in p. 20) corresponds to Fig. 4A in the main text using all bacteria (subplot A), coloured by different features (B-K) or generated using different $\rho_{thr}$ (L-M). Specifically, we can notice that $\rho_{thr}$ = 0.6 provides better graph separability as compared to 0.5 or 0.7 (compare Fig. 4A and subplots L, M in Supplementary Figure 15). Medium-size subgraphs contain abundant species (subplots B, C) with complex dynamics (both stationary and non-stationary, seasonal and non-seasonal, and different occurrence percentage). On the other hand, the cloud is built from bacteria that are different to anything else and are probably the most intriguing. It also exhibits rich dynamics but this is more subject dependent. Seasonal bacteria (**I**, J) can be found in both regions but more seasonal species are present in the cloud of female and donorB subjects as compared to male and donorA. Most abundant species (with largest PC1+PC2 loading; subplots G, H) are present in the connected subgraphs but not all subgraphs comprise such species. Many of them can also be found in the cloud. All regions exhibit high taxonomic diversity (E) but one can identify smaller clusters which are homogeneous. According to expectations, rare bacteria clusters are stationary (K). The same holds for abundant clusters but, interestingly, we can also find large connected components dominated by non-stationary species. The region containing large connected subgraphs is approx. 2-2.5 as large (when it comes to number of species; including rare species) as the cloud irrespective of the dataset - this is, however, dependent on the threshold put on $|\rho|$.

**A**

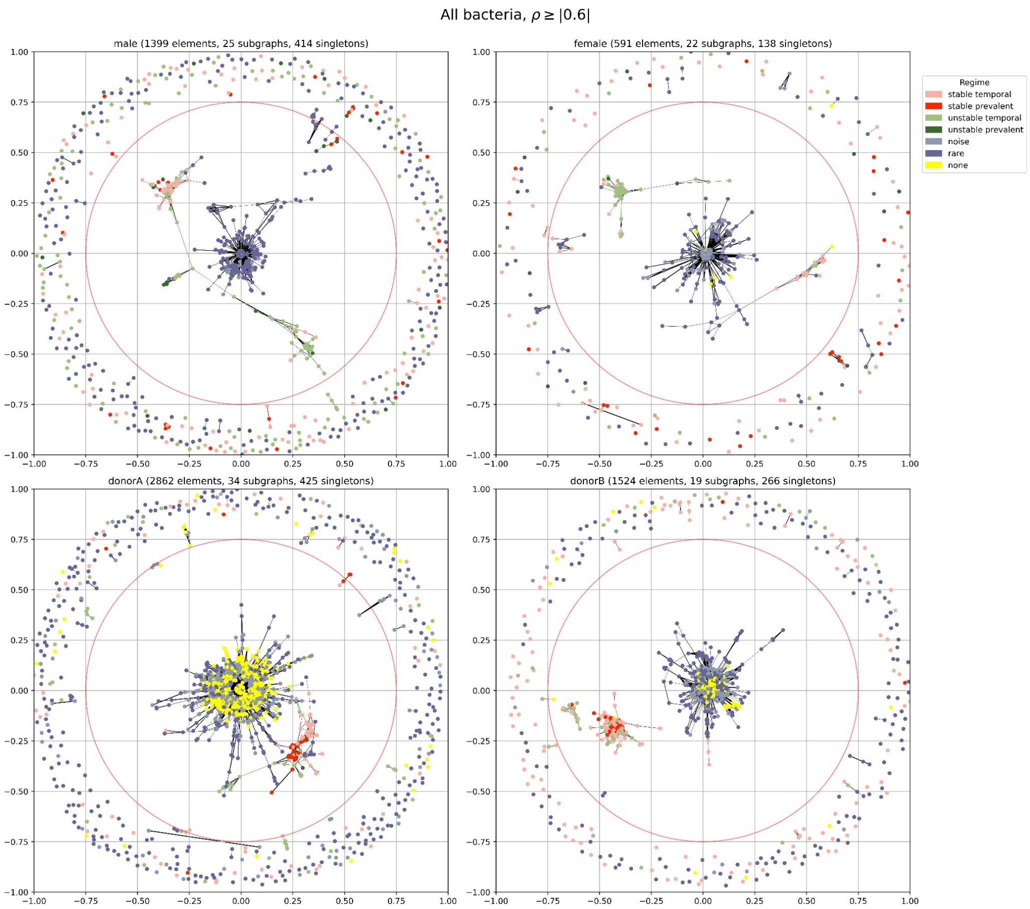

**B**

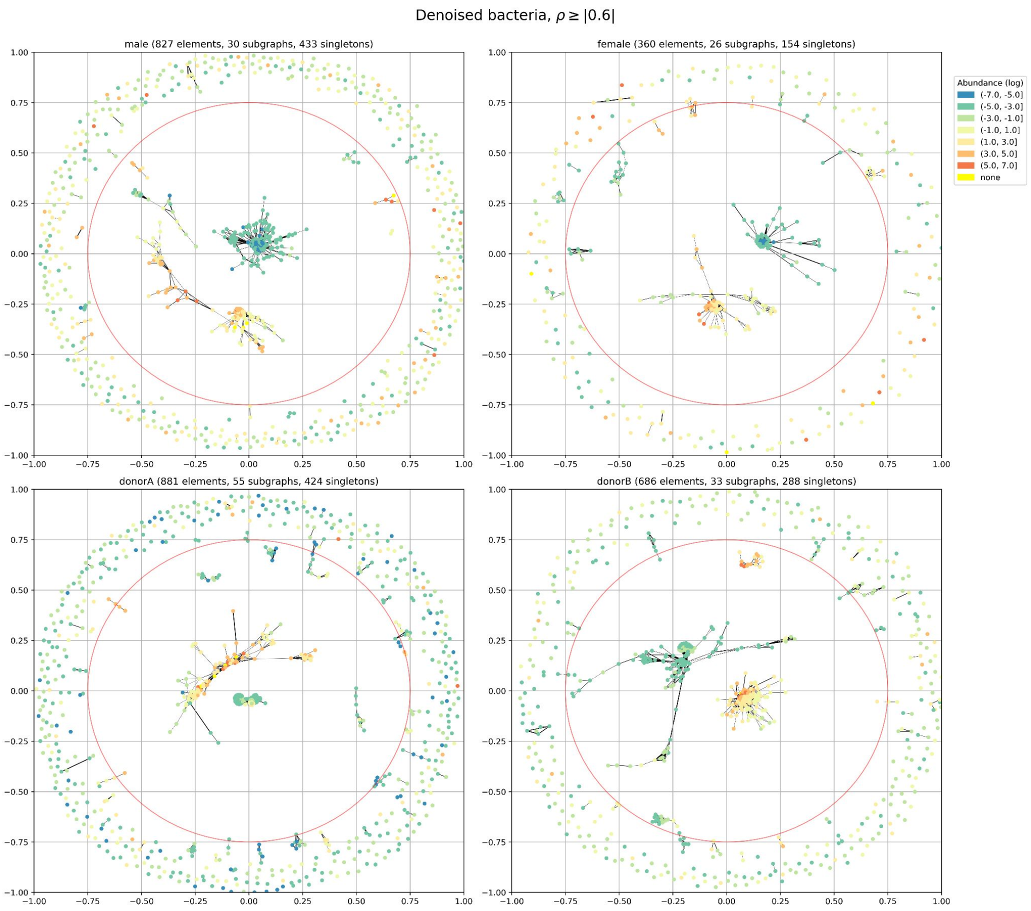
Denoised bacteria, $\rho \geq |0.6|$
male (827 elements, 30 subgraphs, 433 singletons)
female (360 elements, 26 subgraphs, 154 singletons)
donorA (881 elements, 55 subgraphs, 424 singletons)
donorB (686 elements, 33 subgraphs, 288 singletons)
Abundance (log)
(-7.0, -5.0]
(-5.0, -3.0]
(-3.0, -1.0]
(-1.0, 1.0]
(1.0, 3.0]
(3.0, 5.0]
(5.0, 7.0]
none

**C**

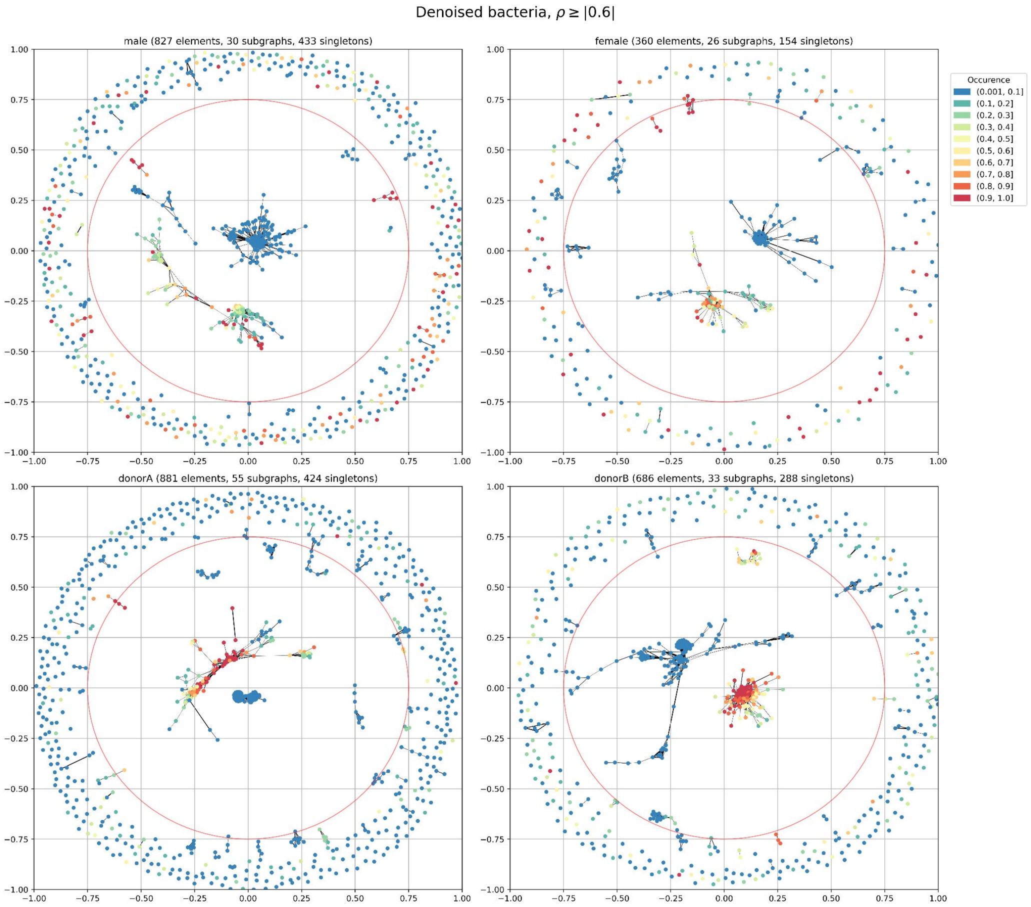
Denoised bacteria, $\rho \geq |0.6|$
male (827 elements, 30 subgraphs, 433 singletons)
female (360 elements, 26 subgraphs, 154 singletons)
donorA (881 elements, 55 subgraphs, 424 singletons)
donorB (686 elements, 33 subgraphs, 288 singletons)
Occurence
(0.001, 0.1]
(0.1, 0.2]
(0.2, 0.3]
(0.3, 0.4]
(0.4, 0.5]
(0.5, 0.6]
(0.6, 0.7]
(0.7, 0.8]
(0.8, 0.9]
(0.9, 1.0]

**D**

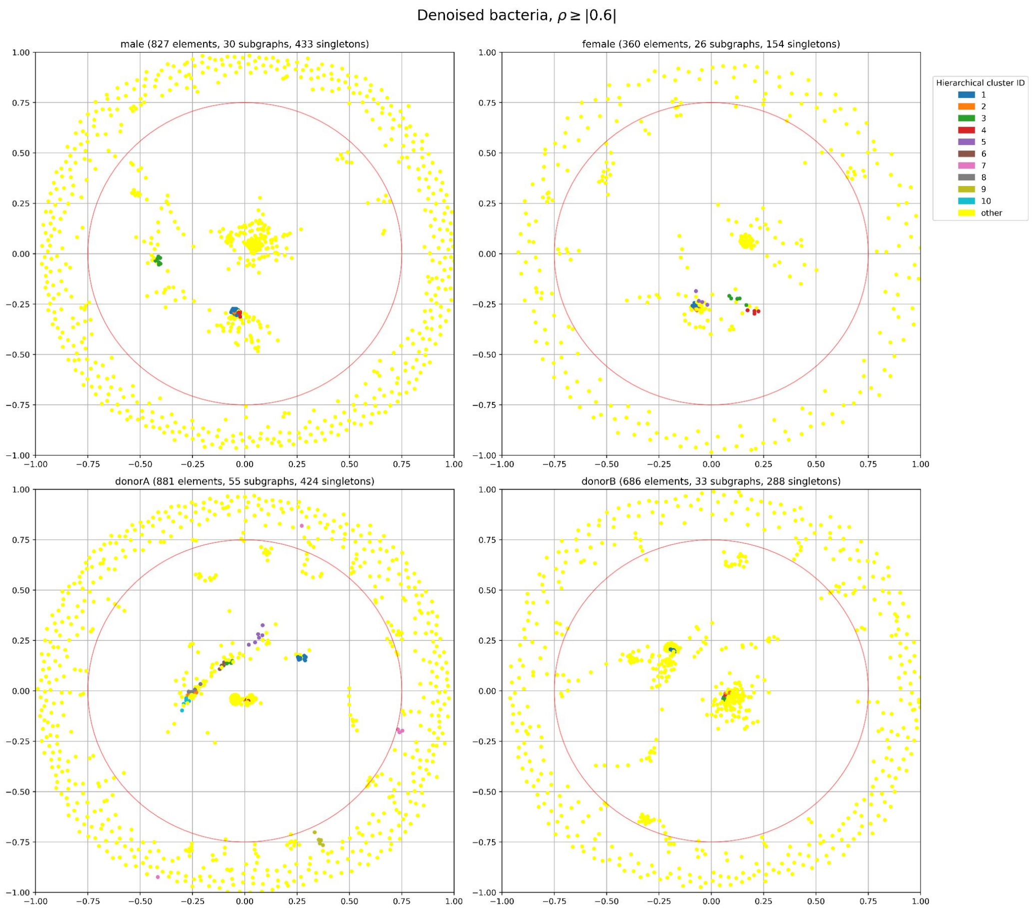
Denoised bacteria, $\rho \geq |0.6|$
male (827 elements, 30 subgraphs, 433 singletons)
female (360 elements, 26 subgraphs, 154 singletons)
donorA (881 elements, 55 subgraphs, 424 singletons)
donorB (686 elements, 33 subgraphs, 288 singletons)
Hierarchical cluster ID
1
2
3
4
5
6
7
8
9
10
other

**E**

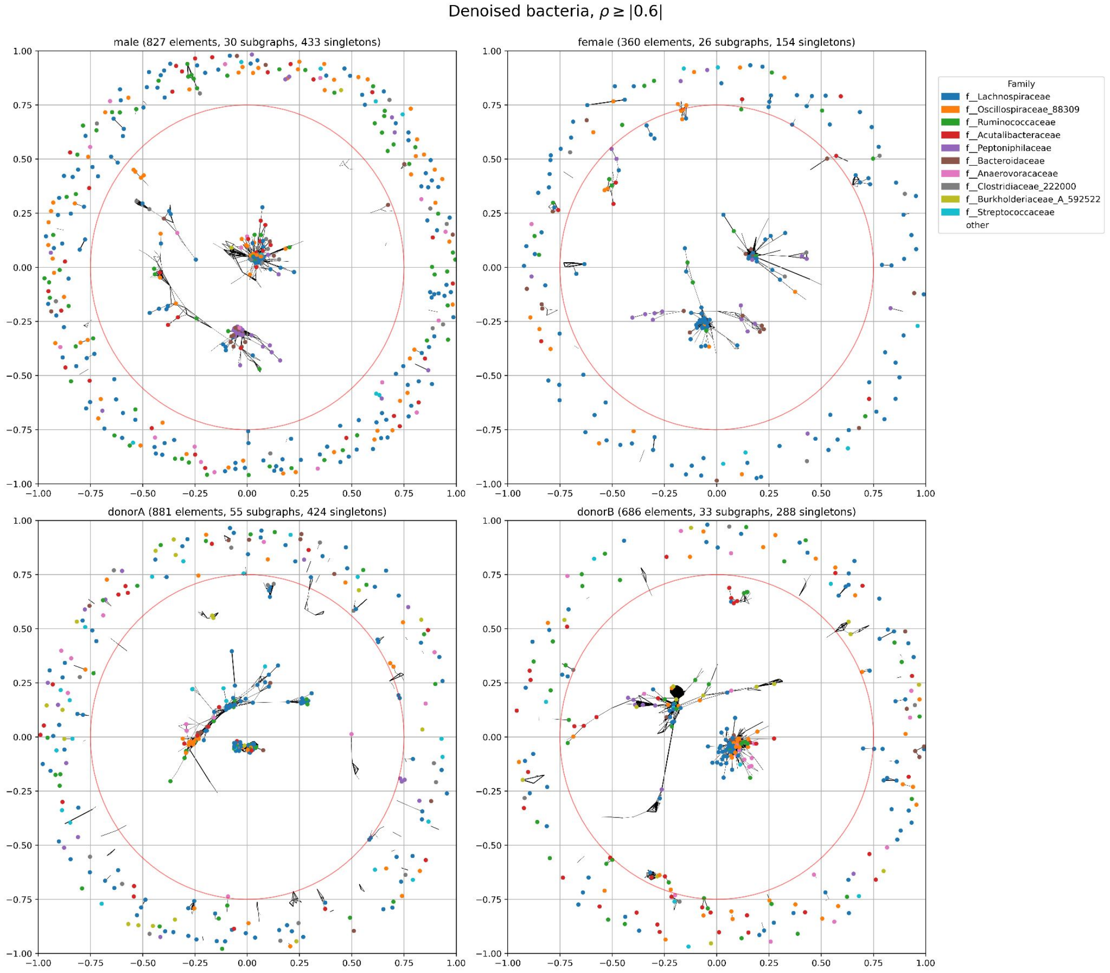
Denoised bacteria, $\rho \geq |0.6|$
male (827 elements, 30 subgraphs, 433 singletons)
female (360 elements, 26 subgraphs, 154 singletons)
donorA (881 elements, 55 subgraphs, 424 singletons)
donorB (686 elements, 33 subgraphs, 288 singletons)
Family
f__Lachnospiraceae
f__Oscillospiraceae_88309
f__Ruminococcaceae
f__Acutalibacteraceae
f__Peptoniphilaceae
f__Bacteroidaceae
f__Anaerovoracaceae
f__Clostridiaceae_222000
f__Burkholderiaceae_A_592522
f__Streptococcaceae
other

F

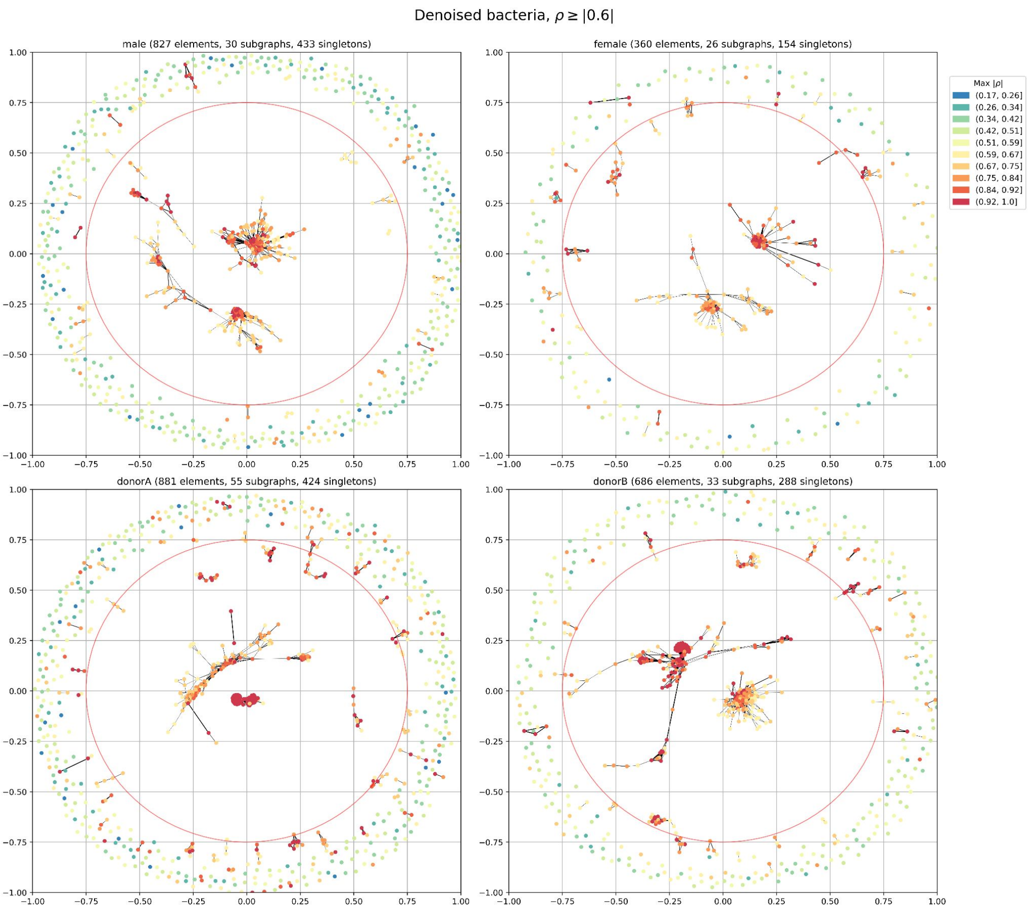
Denoised bacteria, $\rho \geq |0.6|$
male (827 elements, 30 subgraphs, 433 singletons)
female (360 elements, 26 subgraphs, 154 singletons)
donorA (881 elements, 55 subgraphs, 424 singletons)
donorB (686 elements, 33 subgraphs, 288 singletons)
Max $|\rho|$
(0.17, 0.26]
(0.26, 0.34]
(0.34, 0.42]
(0.42, 0.51]
(0.51, 0.59]
(0.59, 0.67]
(0.67, 0.75]
(0.75, 0.84]
(0.84, 0.92]
(0.92, 1.0]

G

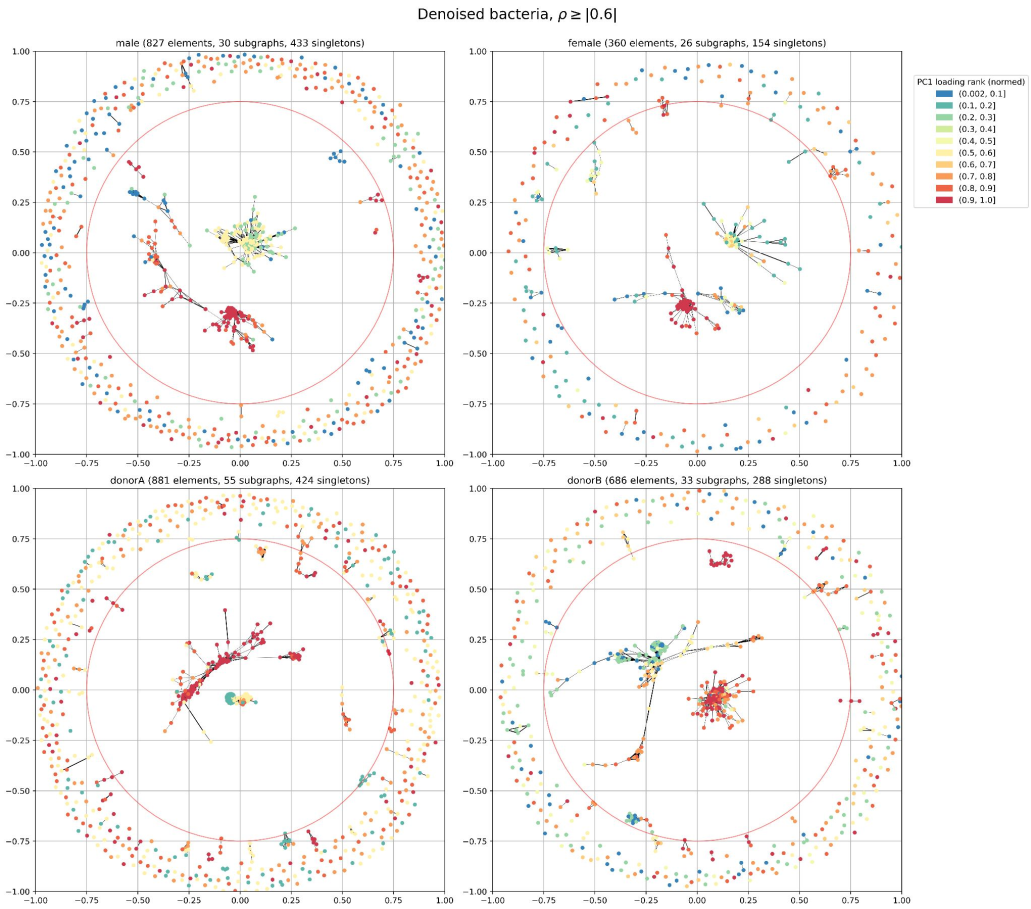
Denoised bacteria, $\rho \geq |0.6|$
male (827 elements, 30 subgraphs, 433 singletons)
female (360 elements, 26 subgraphs, 154 singletons)
donorA (881 elements, 55 subgraphs, 424 singletons)
donorB (686 elements, 33 subgraphs, 288 singletons)
PC1 loading rank (normed)
(0.002, 0.1]
(0.1, 0.2]
(0.2, 0.3]
(0.3, 0.4]
(0.4, 0.5]
(0.5, 0.6]
(0.6, 0.7]
(0.7, 0.8]
(0.8, 0.9]
(0.9, 1.0]

**H**

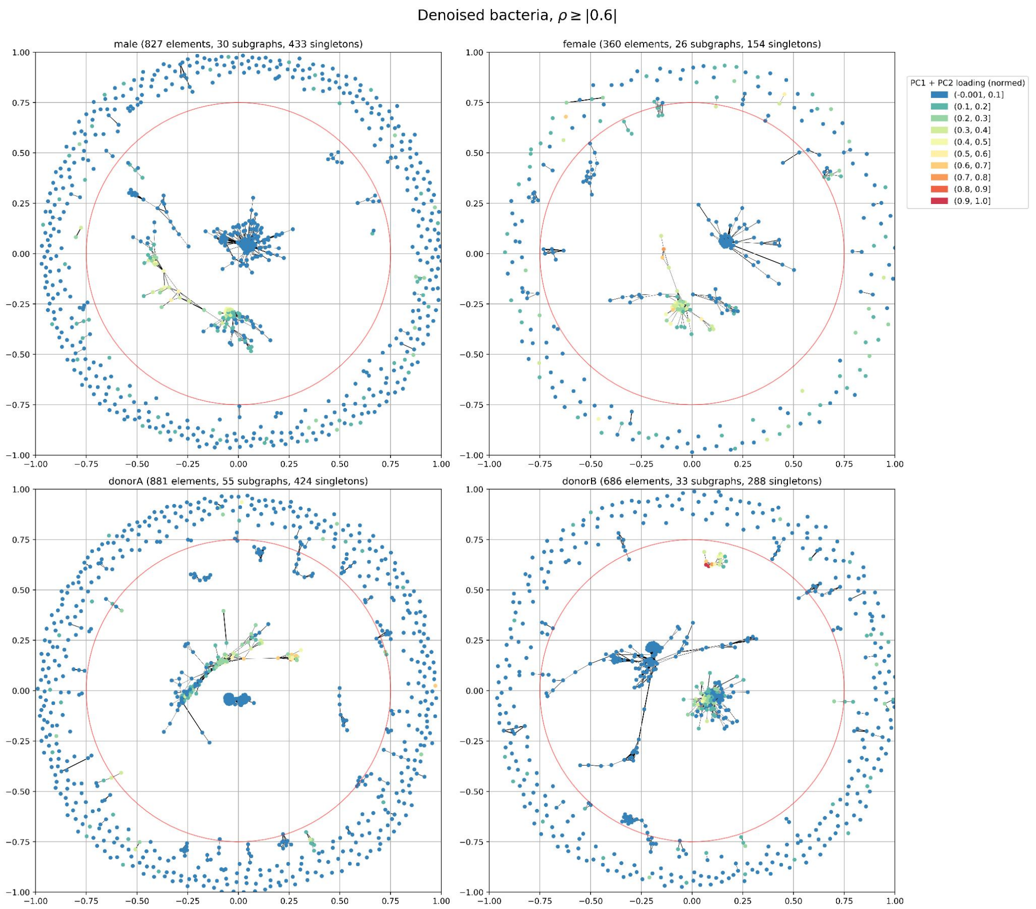

Denoised bacteria, $\rho \geq |0.6|$
male (827 elements, 30 subgraphs, 433 singletons)
female (360 elements, 26 subgraphs, 154 singletons)
donorA (881 elements, 55 subgraphs, 424 singletons)
donorB (686 elements, 33 subgraphs, 288 singletons)
PC1 + PC2 loading (normed)
(-0.001, 0.1]
(0.1, 0.2]
(0.2, 0.3]
(0.3, 0.4]
(0.4, 0.5]
(0.5, 0.6]
(0.6, 0.7]
(0.7, 0.8]
(0.8, 0.9]
(0.9, 1.0]


**I**

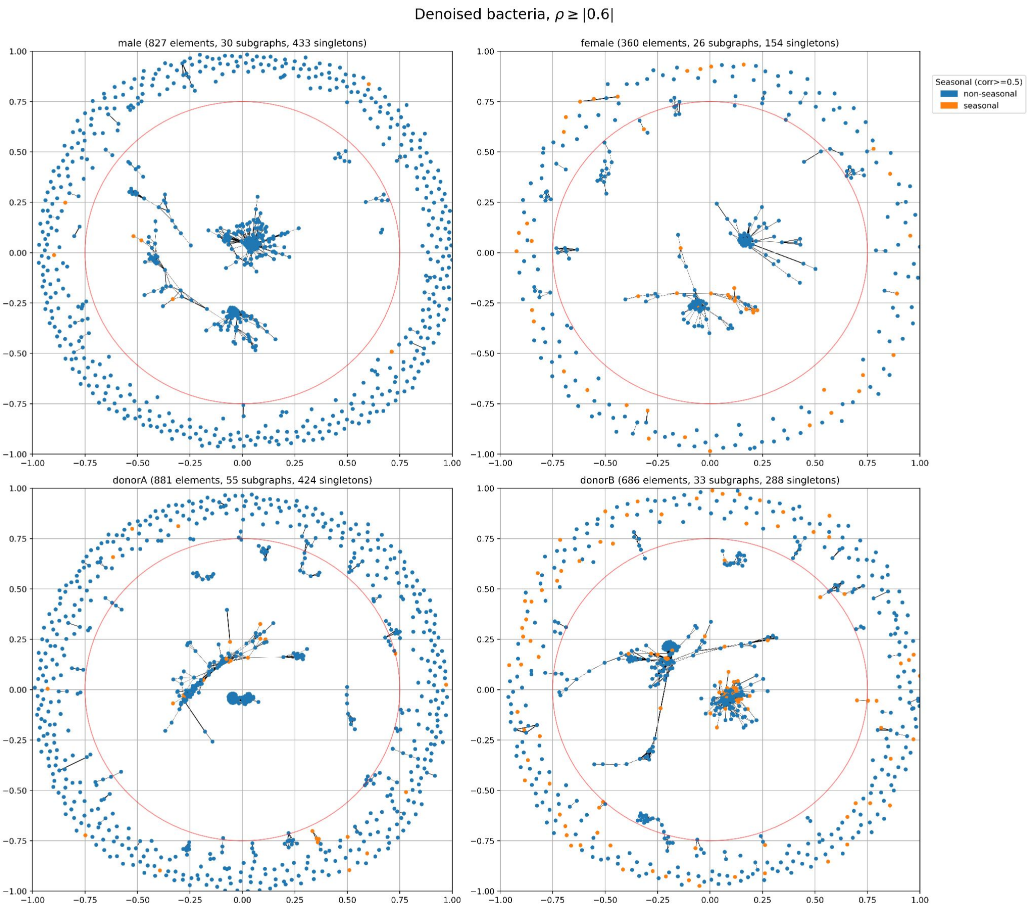

Denoised bacteria, $\rho \geq |0.6|$
male (827 elements, 30 subgraphs, 433 singletons)
female (360 elements, 26 subgraphs, 154 singletons)
donorA (881 elements, 55 subgraphs, 424 singletons)
donorB (686 elements, 33 subgraphs, 288 singletons)
Seasonal (corr>=0.5)
non-seasonal
seasonal

**J**

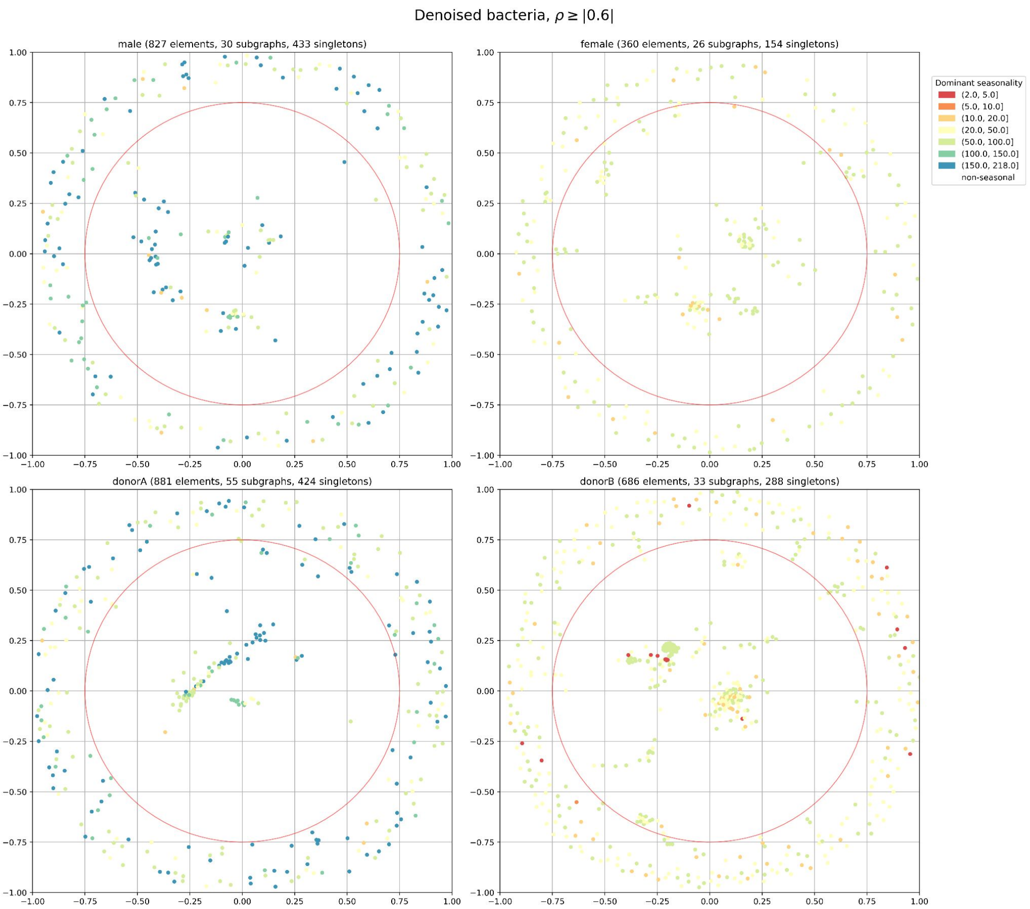
Denoised bacteria, $\rho \geq |0.6|$
male (827 elements, 30 subgraphs, 433 singletons)
female (360 elements, 26 subgraphs, 154 singletons)
donorA (881 elements, 55 subgraphs, 424 singletons)
donorB (686 elements, 33 subgraphs, 288 singletons)
Dominant seasonality
(2.0, 5.0]
(5.0, 10.0]
(10.0, 20.0]
(20.0, 50.0]
(50.0, 100.0]
(100.0, 150.0]
(150.0, 218.0]
non-seasonal

**K**

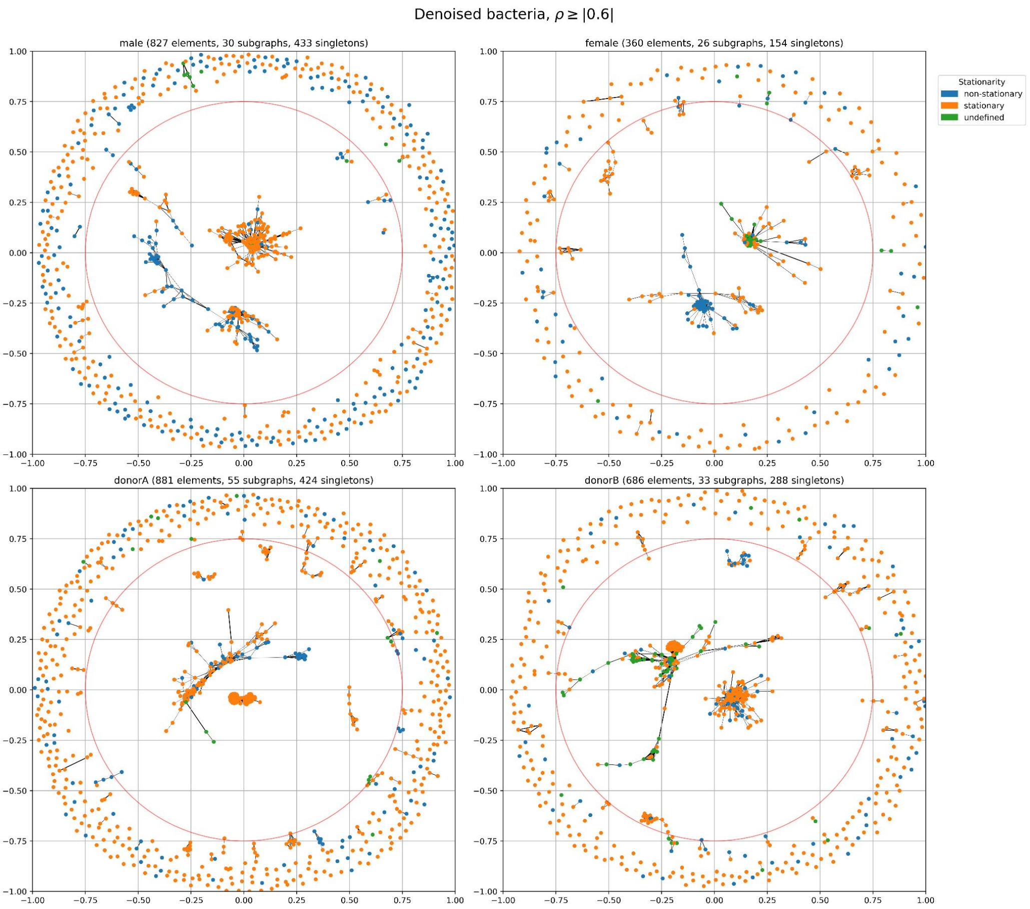
Denoised bacteria, $\rho \geq |0.6|$
male (827 elements, 30 subgraphs, 433 singletons)
female (360 elements, 26 subgraphs, 154 singletons)
donorA (881 elements, 55 subgraphs, 424 singletons)
donorB (686 elements, 33 subgraphs, 288 singletons)
Stationarity
non-stationary
stationary
undefined

L

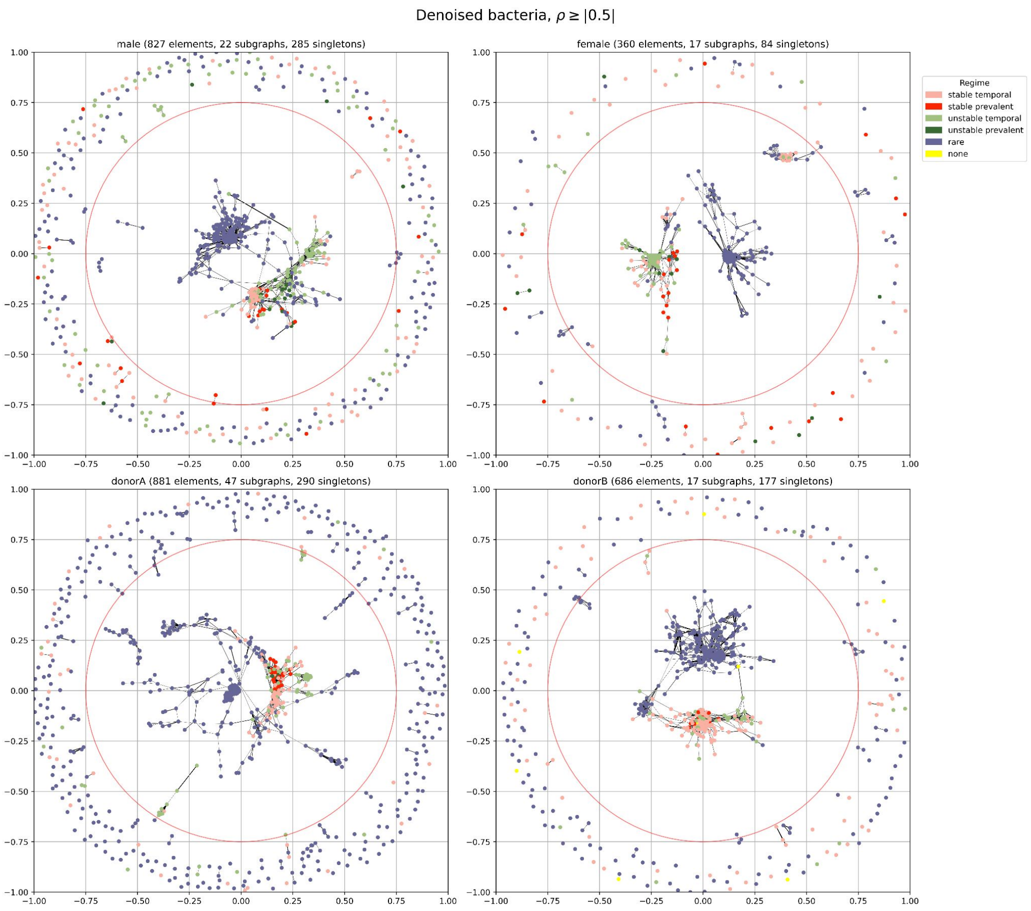
Denoised bacteria, $\rho \geq |0.5|$
male (827 elements, 22 subgraphs, 285 singletons)
female (360 elements, 17 subgraphs, 84 singletons)
donorA (881 elements, 47 subgraphs, 290 singletons)
donorB (686 elements, 17 subgraphs, 177 singletons)
Regime
stable temporal
stable prevalent
unstable temporal
unstable prevalent
rare
none

M

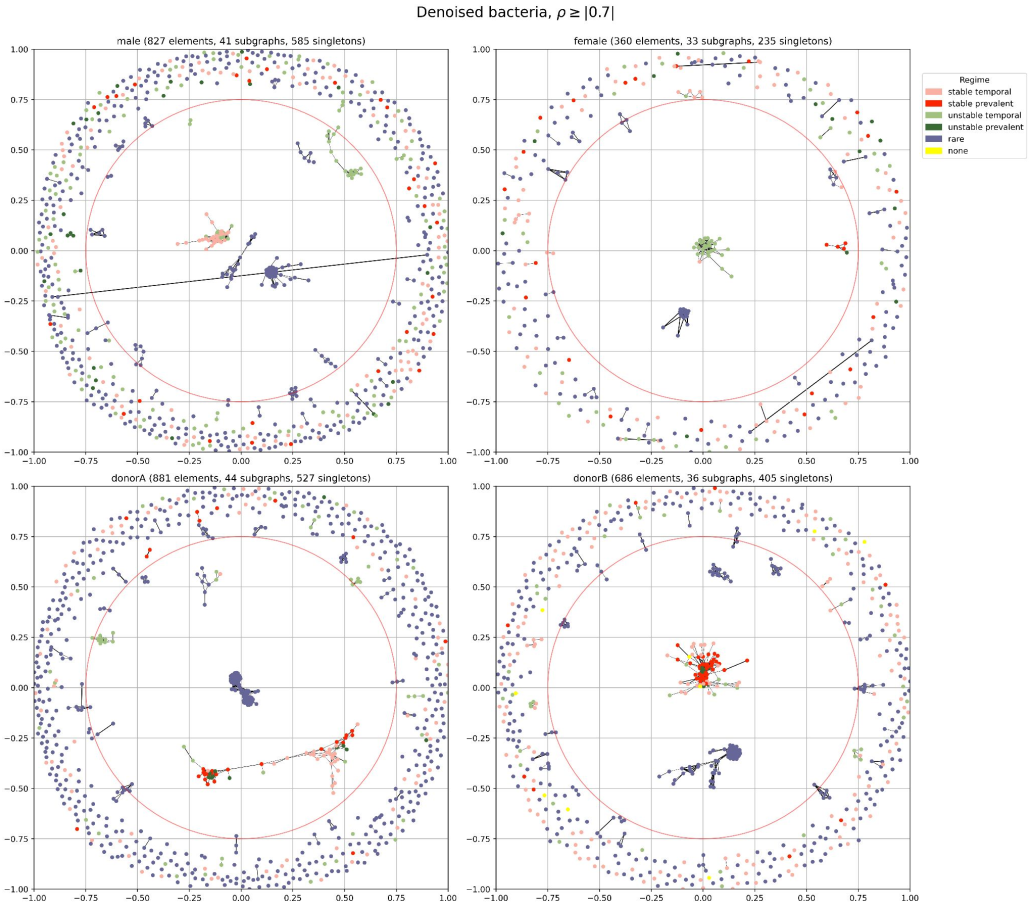
Denoised bacteria, $\rho \geq |0.7|$
male (827 elements, 41 subgraphs, 585 singletons)
female (360 elements, 33 subgraphs, 235 singletons)
donorA (881 elements, 44 subgraphs, 527 singletons)
donorB (686 elements, 36 subgraphs, 405 singletons)
Regime
stable temporal
stable prevalent
unstable temporal
unstable prevalent
rare
none

**Supplementary Figure 15. Results of cluster analysis performed with NetworX.** Each subplot (**A-M**) represents a network (one panel per subject) of bacterial species (nodes) colored by different feature, where edges represent connections equal or stronger than $\rho_{thr}$ = 0.6 (equivalent to $|\rho| \geq 0.6$) except subplot L and M, where $\rho_{thr}$ = 0.5 and 0.7 have been used. In subplots A, B, L and M "none" represents bacteria that didn't pass rarefaction. Red circle in each panel separates the inner part from the cloud.

### Regime evolution in time

Supplementary Figure 16 (see caption in p. 22) corresponds to Fig. 4B-C but for female and donorA subjects (panels A and B) or for male subject but generated using different $\rho_{thr}$ (panels C, and D).

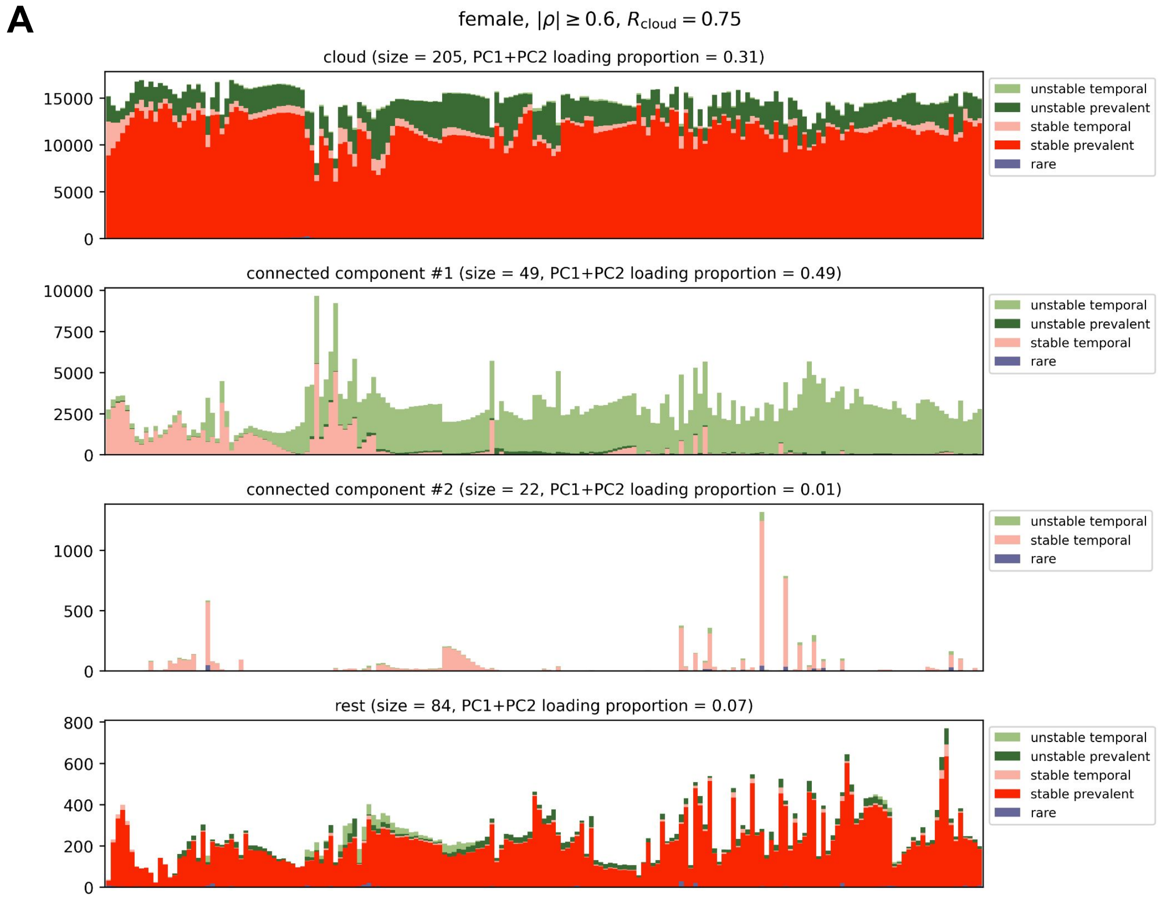

B

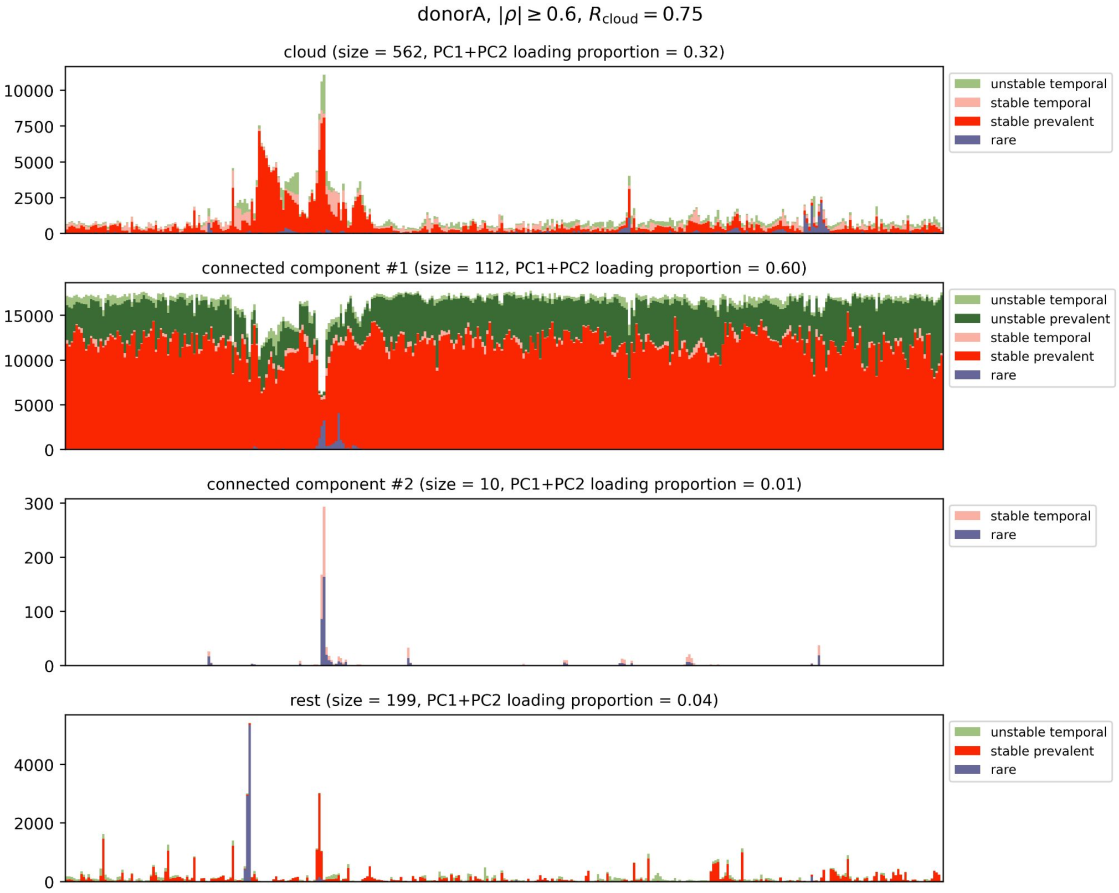
donorA, $|\rho| \geq 0.6$, $R_{\mathrm{cloud}} = 0.75$
cloud (size = 562, PC1+PC2 loading proportion = 0.32)
unstable temporal
stable temporal
stable prevalent
rare
connected component #1 (size = 112, PC1+PC2 loading proportion = 0.60)
unstable temporal
unstable prevalent
stable temporal
stable prevalent
rare
connected component #2 (size = 10, PC1+PC2 loading proportion = 0.01)
stable temporal
rare
rest (size = 199, PC1+PC2 loading proportion = 0.04)
unstable temporal
stable prevalent
rare

C

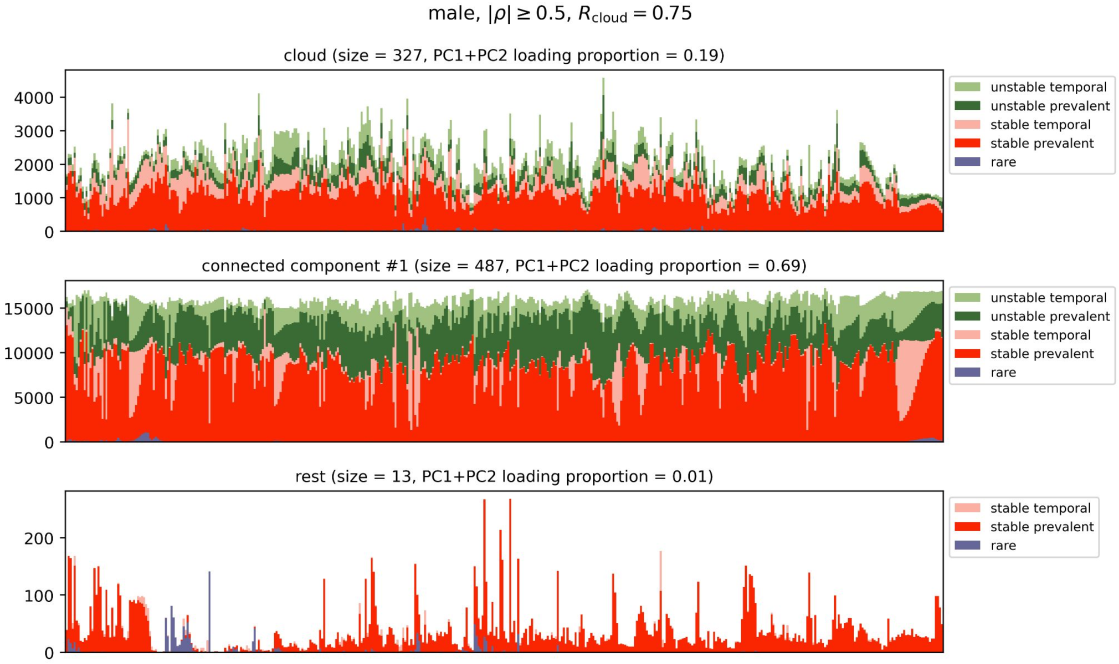
male, $|\rho| \geq 0.5$, $R_{\mathrm{cloud}} = 0.75$
cloud (size = 327, PC1+PC2 loading proportion = 0.19)
unstable temporal
unstable prevalent
stable temporal
stable prevalent
rare
connected component #1 (size = 487, PC1+PC2 loading proportion = 0.69)
unstable temporal
unstable prevalent
stable temporal
stable prevalent
rare
rest (size = 13, PC1+PC2 loading proportion = 0.01)
stable temporal
stable prevalent
rare

**D**

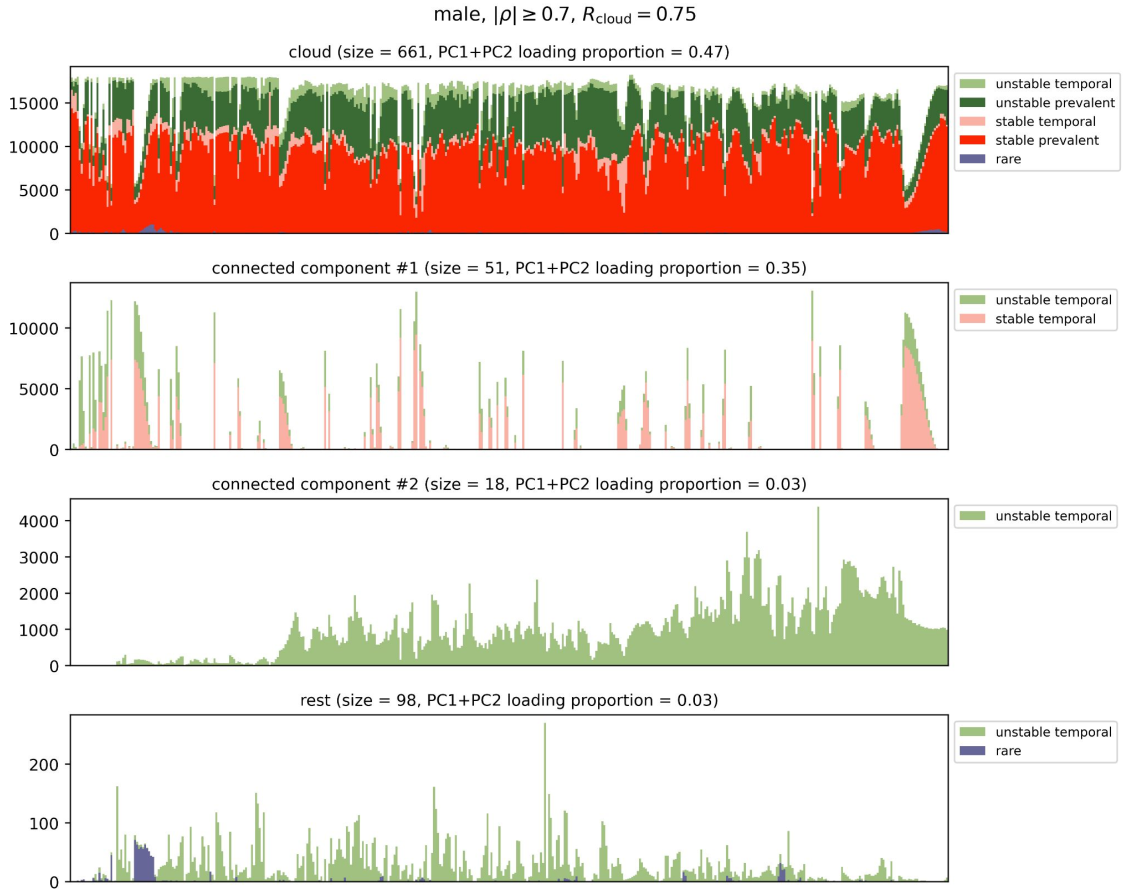


**Supplementary Figure 16. Change in total counts after rarefaction over time** for female (**A**), donorA (**B**) and male (**C, D**) subject stratified by regime (color) and group of bacteria (panels) i.e. cloud, largest connected components and the rest. In panel A and B $\rho_{thr}$ = 0.6 whereas in panel C and D $\rho_{thr}$ = 0.5 and 0.7 respectively. $R_{cloud}$ represents a diameter that separates the inner part from the cloud.

### Examples

Supplementary Figure 17 shows same example bacteria clustered together in the largest connected components. Heatmaps showing their mutual proportionality |ρ| are presented in Supplementary Fig. 18. Clearly, species placed close to each other on the graph have larger proportionality, which stays in line with the analysis presented in the next section (see Supplementary Fig. 19). Interestingly, in same cases (bacteria 8 and 9 in group A for donorA, bacteria 2 and 3 in group B for donorB) we can notice anti co-occurrence, but in order to shed more light on the underlying effect behind it, both higher taxonomic resolution and functional analysis are required.

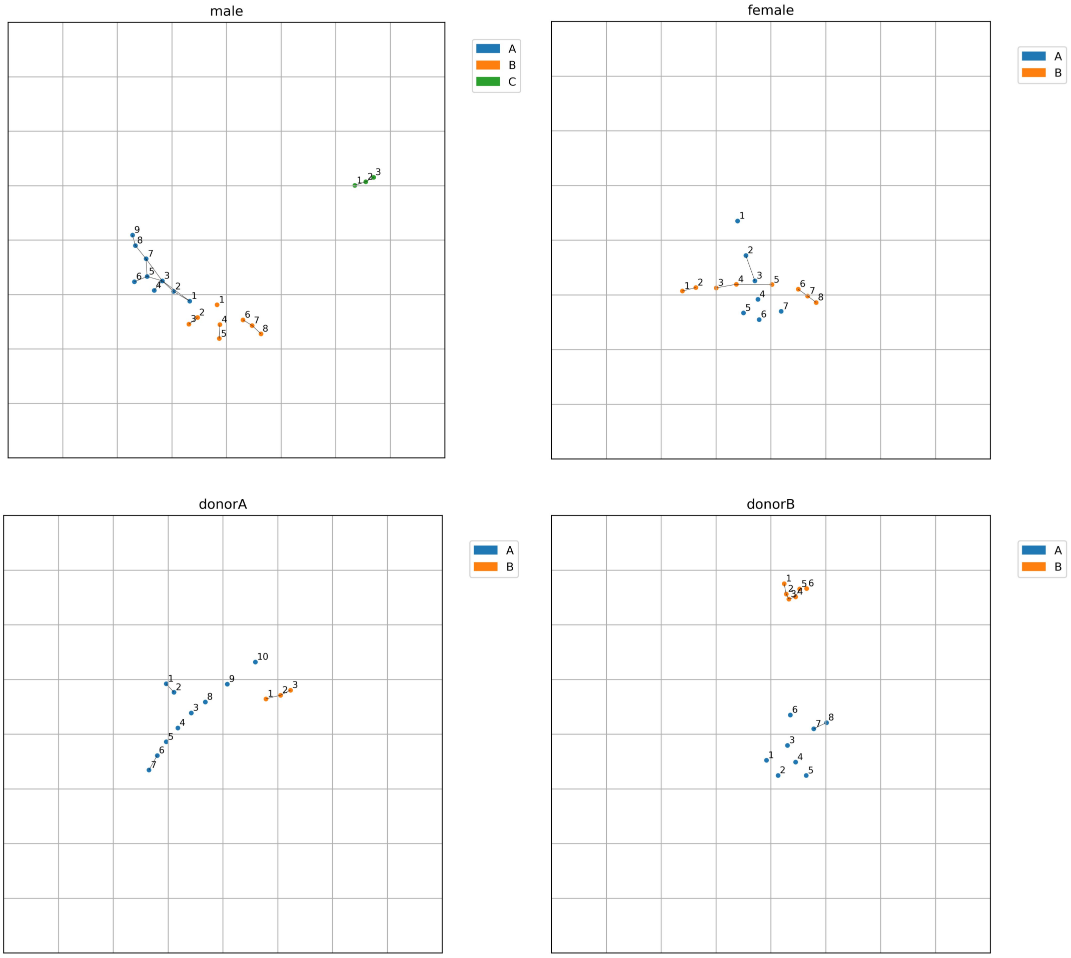


**Supplementary Figure 17.** Example bacteria selected from NetworkX graph shown in Fig. 4A in the manuscript. Each color for every subject represents a different group (cluster) of bacteria.

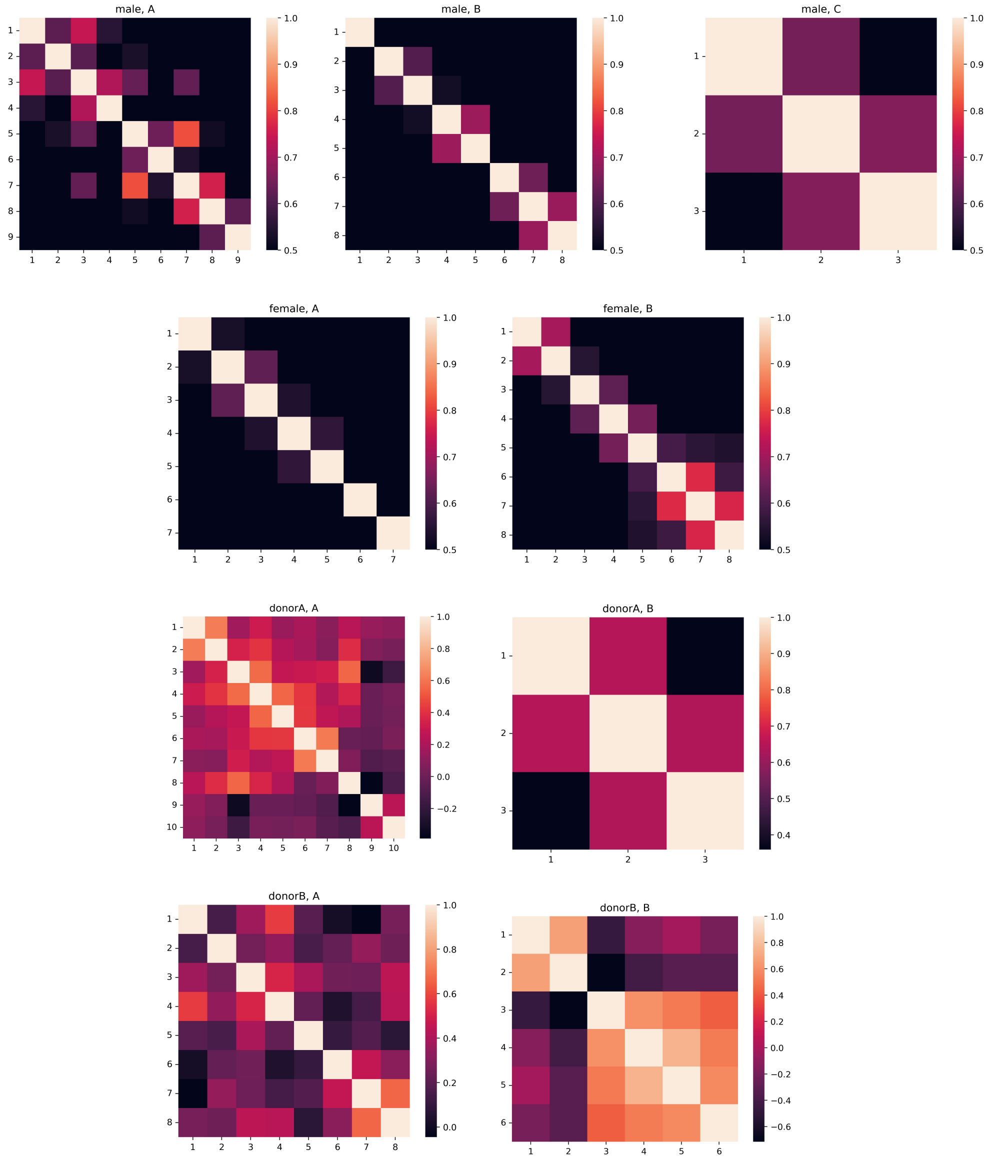


**Supplementary Figure 18.** Heatmaps showing proportionality between bacteria presented in Supplementary Figure 17 (one heatmap per bacterial group).

### Sanity check - Euclidean distance vs $|\rho|$

In order to check whether similar bacteria are placed close to each other in the graphs we compared absolute value of their proportionality $|\rho|$ against Euclidean distance (Supplementary Figure 19). Indeed, we can see clear relation i.e. the larger $|\rho|$ the smaller distance which is evident especially for the largest connected components (left panels). When it comes to the cloud (right panels) existence of few points above the threshold $\rho_{thr}$ (=0.6) comes from the fact that the cloud definition is imperfect and some smaller clusters were tiered apart by imposing the criterion on the cloud diameter value ($R_{cloud}$ = 0.75). Distance distribution for $|\rho| \leq 0.6$ on the right panels shows that the polar angle doesn't matter for distant points (singletons in the cloud) - an extreme example is visible in the top left panel in Supplementary Figure 15M. However, this conclusion doesn't hold for the inner regions which are of our special interest.

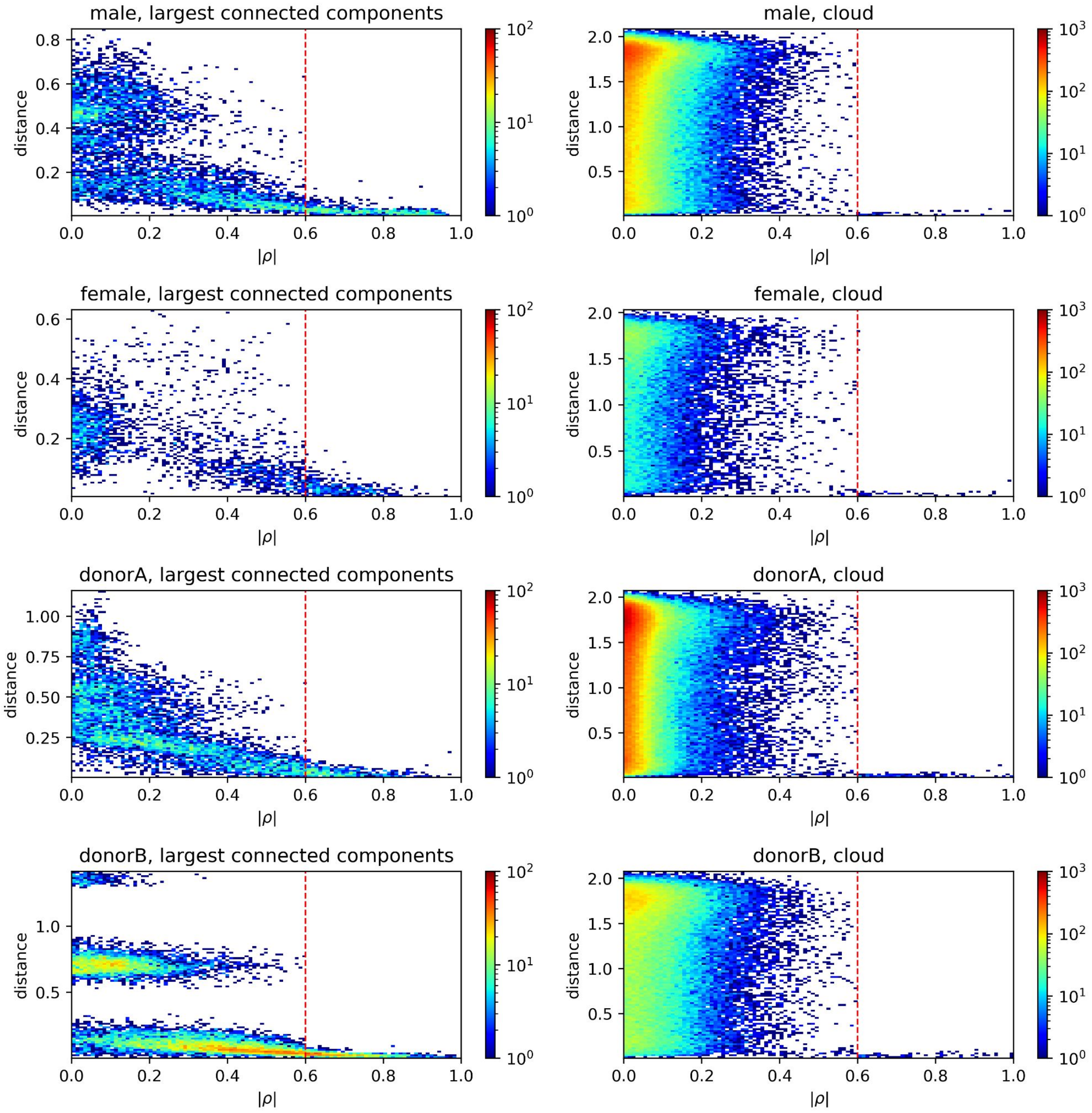


**Supplementary Figure 19:** Relationship between proportionality |$\rho$| and Euclidean distance of bacteria shown in NetworkX graph in Fig. 5 in the manuscript.

### Hierarchical clustering

Using *D* as a feature matrix (treating it as a usual distance matrix didn't perform well) we constructed hierarchical clusters in order to compare them with the NetworkX graphs. As a sanity check we computed intra-cluster proportionality. According to expectations, the proportionality is high especially for the most abundant ones (see Supplementary Figure 20 and 21). Panel D in Supplementary Figure 15 shows location of the most abundant clusters (with cardinality ≥ 5 and mean abundance after rarefaction ≥ 0.1). Interestingly, they are mainly located in the central region of the graph (comprising largest connected components) supporting the hypothesis that this is the main part of the graph that drives the microbiome dynamics. We can also find that some clusters e.g. 1, 2 and 4 for male subjects are placed very close to each other. Indeed, in this example the inter-cluster similarity is very high ($\rho$ = 0.79 ± 0.12) and they can be treated as one larger cluster. It demonstrates that the NetworkX approach is more robust although it doesn't allow for strict cluster extraction.

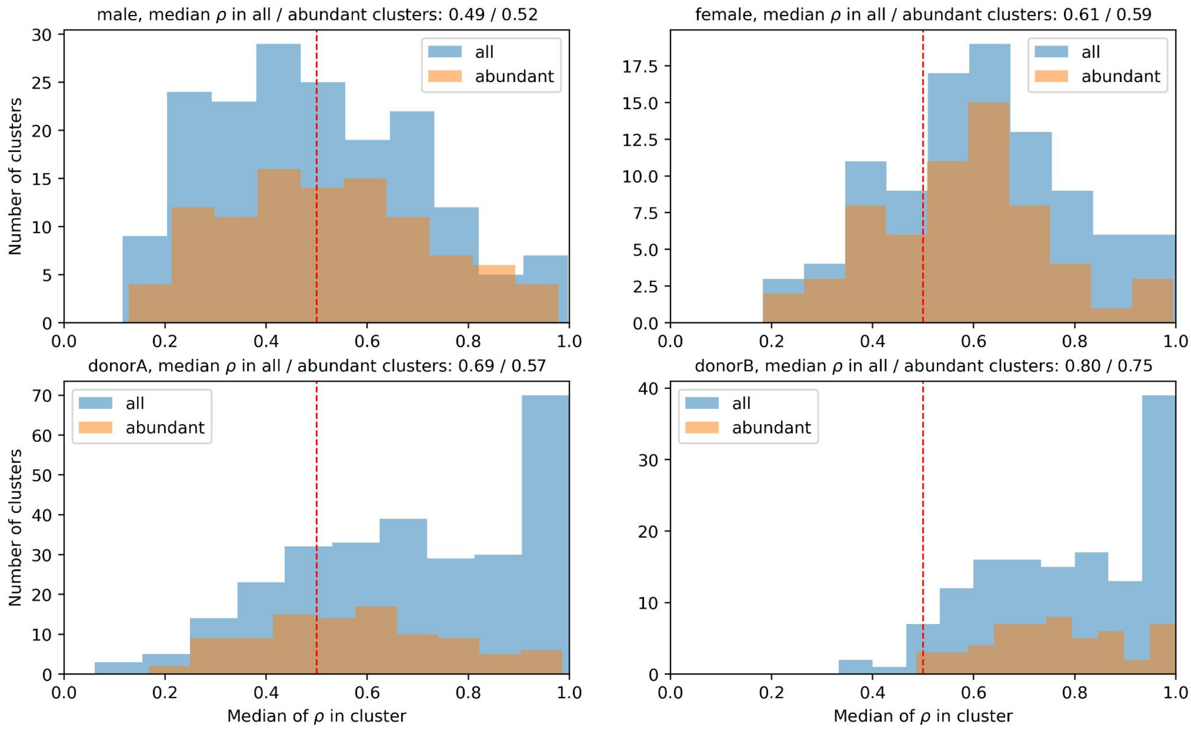


**Supplementary Figure 20. Results of hierarchical clustering.** Distribution of intra-cluster proportionality. Abundant clusters are clusters with mean bacterial abundance equal or higher than 0.1.

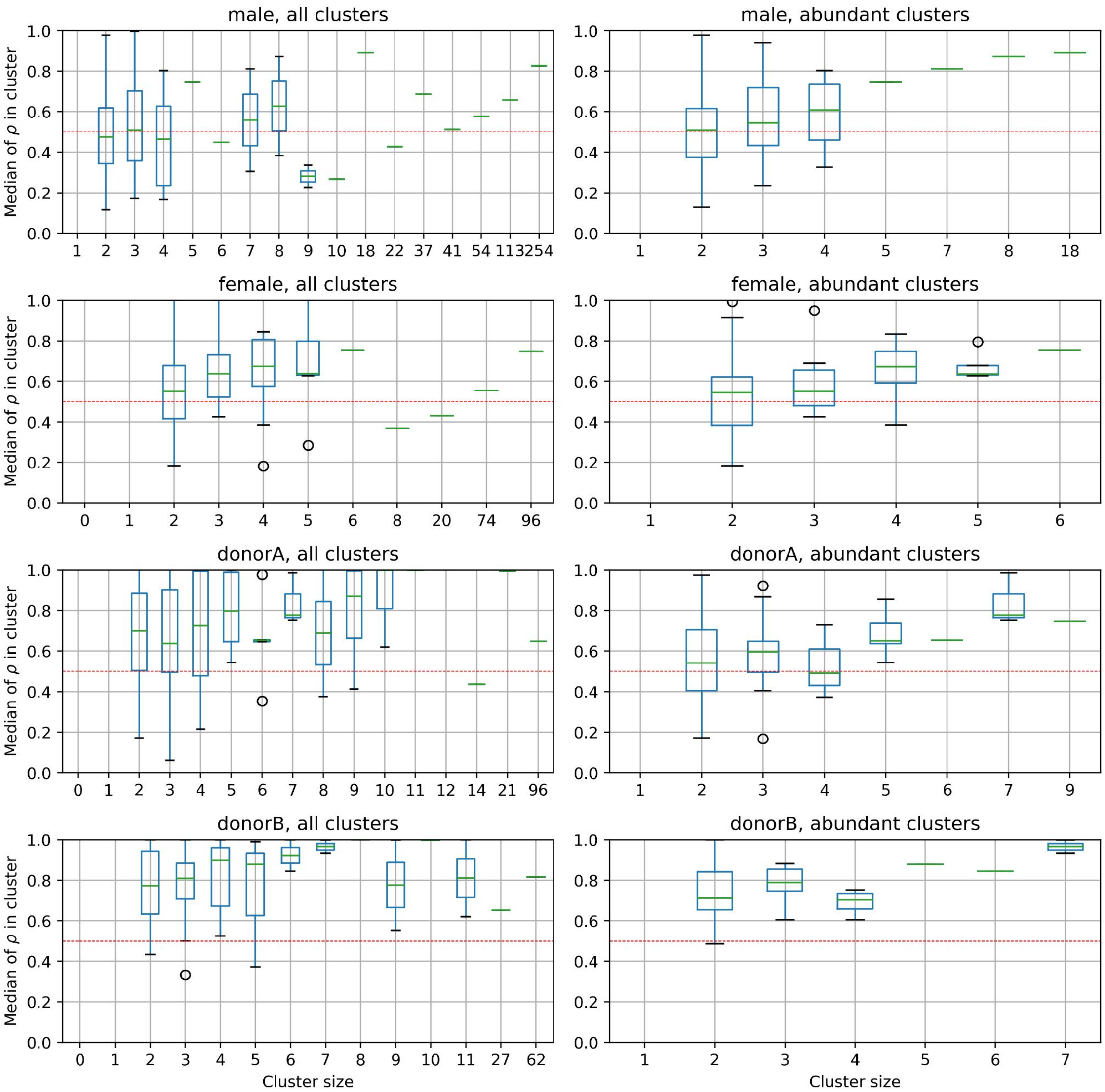


**Supplementary Figure 21.** The same as in Supplementary Figure 20 but each box plot corresponds to mean $\rho$ distribution of bacteria in clusters of a given size (shown in x-axis).
